# Differential expression of keratin and keratin associated proteins are linked with hair loss condition in spontaneously mutated inbred mice

**DOI:** 10.1101/2024.07.13.603037

**Authors:** Neeraja Chilukoti, Sivapriya Pavuluri, Satish Kumar

## Abstract

Hair loss condition is heritable and is influenced by multifactorial inheritance. In the present study, spontaneously mutated mice showed hair loss phenotype with defect in the first cycle of hair follicle formation leading to cyclic alopecia. These mutant mice follow autosomal recessive inheritance pattern. The transcriptomic profile and differential gene expression analysis of skin tissues by RNA-sequencing at different stages of hair cycle formation was performed. The genes with significant differential genes expression levels in each stage of hair cycle formation were identified and most of these genes were shown to be associated with keratinization process and hair follicle formation. Transcriptome profiling followed by QPCR validation revealed that mRNA levels of *Krt16*, *Alox15*, *Fetub* (upregulated) and *Msx2* (downregulated) were significantly differentially expressed in mutant skin tissues during late anagen and catagen stages. *Krt6b* mRNA and protein levels were significantly higher in the mutant mice during all stages of first hair cycle formation. The present study provides basis for understanding the differential gene expression of hair-related genes, including keratinization-associated proteins and its relevance. These mutant mice can serve as a model for studying hair loss condition that can be further used in the identification, evaluation and treatment strategies for alopecia condition.

## 1. Introduction

Hair follicles are mini appendages of skin that consists of outer root sheath in continuation with the epidermis, inner root sheath and hair shaft. Hair follicles continuously undergo different stages of growth (anagen), regression (catagen) and rest (telogen) (Schneider *et al*. 2009). Several genes along with the surrounding epidermal and dermal layer specific keratin and keratin associated proteins are responsible for promoting the hair follicle growth (Cai *et al*. 2009). Keratins are intermediate filaments that are also expressed in the hair follicles along with hair stem cells essential for growth and development (Langbein *et al*. 2001). These keratin proteins form assembled strong networks that help keratinocytes attach together and connect the epidermis to the underlying skin layers contributing to the structure and growth of the hair follicle (Schweizer *et al*. 2007). Abnormal keratinization leads to hair loss condition due to defect in the genes contributing to different types of keratin expression that are of importance in hair follicle formation (Oka *et al*. 2020). However, the genes and molecular mechanisms involved in the abnormal hair growth leading to hair loss have not yet been elucidated.

The hair follicles undergo changes in their shape during their different growth stages (anagen, catagen and telogen). These growth changes during different phases are closely regulated and connected with the activation and regulation of a range of signaling pathways. Various signaling pathways such as Wnt/β-catenin, sonic hedgehog (Shh), and notch pathways were known to be involved in the entry of hair follicle to anagen phase. Wnt/β-catenin pathway plays an important role in the in the transition of phase from telogen to anagen (Zhang *et al*. 2020). Earlier studies revealed several molecular mechanisms that control the hair follicle and epidermal homeostasis (Kobielak *et al*. 2007; Eckhart *et al*. 2013; Lim and Nusse 2013). These studies and other several transcriptomic analyses found genes especially expressed in different cell populations of the hair follicles and epidermis that code for transcription factors regulating gene expression in the skin (Joost *et al*. 2016; Yang *et al*. 2017; Cheng *et al*. 2018). Most of these data lack information on growth phase specific gene expression that may provide essential information on genes and pathways that are required for hair follicle formation and homeostasis.

In the present study, we investigated the transcriptomic profile and differential gene expression of skin tissues belonging to wildtype and spontaneously mutated C57BL/6 mice by RNA sequencing at different stages of first hair cycle formation. The present study provides insights into the genes that are specifically expressed in anagen, catagen and telogen of hair cycle formation in the spontaneously mutated mice. Also, these mutant mice can serve as model organism and aid in the development of novel strategies to control the hair loss condition.

## 2. Materials and methods

### 2.1 Animals

The present study was accepted by Institutional Animal Ethics Committee, Centre for Cellular and Molecular Biology, Hyderabad, India (Animal trial registration number 20/1999/CPCSEA dated 10/3/99). All mice studies were executed according to the approved institutional ethical guidelines (IAEC76/ CCMB/ 2019) of Centre for Cellular and Molecular Biology, Hyderabad, India. C57BL6J mice were housed in protected environment with 12 h light–dark cycle and were fed ad libitum.

### 2.2 Mice Inbreeding

Male mice with hairloss (HL) phenotype were mated with wild type (WT) females. The F1 progeny was heterozygous (HT) with hairy skin. Subsequently, the litter mating of F1 mice resulted in F2 progeny with a ratio of 3:1 corresponding to WT/HT to HL. To identify WT from HT mice, mice were backcrossed with the F1 HT mice. The HL mice were continuously inbred to maintain the genetic factors contributing to the HL phenotype.

### 2.3 Morphological studies

HL mice were continuously observed for hairloss patterns at different postnatal days (P0, P16, P19 and P28) corresponding to different stages of hair cycle formation (anagen, catagen, telogen and anagen) respectively.

### 2.4 Growth and fertility of HL mice

Postnatal male pups of WT, wild type colony (WTC) and HL animals were monitored for body weight and body length at P0, P16, P19, P28 stages. Fertility of HL animals was assessed by setting up the crosses: HL male x HL female, WTC male x WTC female, HL male x WTC female, WTC male x HL female. The pups from the above mentioned crosses were monitored for body weight, body length and litter size.

### 2.5 Histological analysis of HL and WT mice

Skin was excised from the dorsal side of the male mice with genotype WT and HL. One cm skin sections corresponding to different stages of hair cycle formation at different postnatal days (P0, P16, P19, P28). Collected skin sections were fixed in 10% formalin, embedded in paraffin wax for histological examination. Thin sections were made out of skin samples (4 μm) and mounted on positively charged probe on plus slides (Fischer scientific, USA). Hematoxylin and eosin staining of skin sections were performed and images of these sections were captured at 100X magnification.

### 2.6 Transcriptome sequencing and data analysis

Total RNA was extracted from the dorsal skin of HL and WT male mice using RNeasy Kit (Qiagen). RNA samples corresponding to postnatal days, P0, P16, P19, P28 were collected in duplicates. TruSeq sample preparation protocol was used to capture transcripts of interest. The transcript products were then purified and further enriched with PCR reaction to create the ultimate cDNA library. The libraries were sequenced in HiSeq X10 for 150bp read length using Sequencing Chemistry v4.0. Quantification of the cDNA library templates and quality control analysis were performed to generate optimum cluster densities throughout every lane of the flow cell. Paired end library was constructed to obtain a sequencing read length of 2×150 bp. Reference based transcriptome sequencing was performed using HiseqX10 platform. Appropriate filters and cut off were set to obtain quality data. From the raw reads, the adapter sequences are removed using cutadapt tool (v-2.6). The expression estimation is performed for all the samples and then it test for the statistical significance of each observed change in expression between the samples. To find differentially expressed genes, it is presumed that the number of reads generated by each transcript is proportional to its large quantity. Cuffdiff platform was used to analyze the differential expression of genes and transcripts, thus facilitating in understanding transcriptional and post transcriptional regulation under different conditions. Further, the results were then annotated from the uniprot and gtf.

### 2.7 Gene ontology and pathway analysis

DAVID annotation tool was used to find significant pathways (KEGG) for the differentially regulated genes. In addition, Integrated Differential Expression and Pathway analysis (iDEP.96 version) was used to find the important biological process and pathways. Expression based heatmap was created using Heatmapper software. The RNA sequencing data of different stages (P0, P16, P19 and P28) of first hair cycle formation generated in the current study was assigned data series number GSE220769 at NCBI portal. The link for the same can be obtained through the link – https://www.ncbi.nlm.nih.gov/geo/query/acc.cgi?acc=GSE220769.

### 2.8 Quantitative PCR

Total RNA was isolated from dorsal skin of WT and HL mice at different postnatal days using RNeasy Kit (Qiagen). Reverse transcription was performed using reverse transcriptase kit (Takara). Quantitative PCR was performed using SYBR Green (Takara, Japan) as described earlier (Singh *et al*. 2020). Ribosomal protein L13 (Rpl13a) was used as a reference gene. Primer sequences used in the present study were listed in supplementary table 1.

### 2.9 Western blotting

Dorsal skin of WT and HL male mice was collected at different postnatal days P0, P16, P19 and P28 and stored at –80°C. Skin tissue was lysed using RIPA buffer and the obtained lysates were quantified for the protein concentration using BCA Protein Assay Kit (Thermo Scientific, USA). 30 μg of proteins were resolved on 10% SDS-PAGE and transferred onto the PVDF membrane. The membrane was then probed with anti-krt6b antibody (Abcam), anti-Msx2 antibody (Abcam), anti-alox15 antibody (Abcam), anti β-actin antibody (Novus Biologicals). Beta actin was used as a loading control for western blotting. HRP-conjugated secondary antibodies were used for chemiluminiscent detection of the proteins. Quantification of protein expression was carried out using Image J software.

### 2.10 Statistical analyses

Statistical analyses were executed using GraphPad Prism 5 software (La Jolla, USA). Data generated were represented as mean ± S.E.M. Unpaired t-test and a two-way ANOVA analyses were performed to define the statistical significance between two groups and multiple groups.

## 3. Results

### 3.1 Autosomal recessive inheritance of hairloss phenotype in C57BL6J mice

Mating of wildtype (WT) female mice with hairloss and mutant (HL) male mice resulted in progeny with 100% WT phenotype. This observation clearly indicates that the F1 progeny is carrying the mutations for HL with no visible variation in hair growth. Litter mating of F1 mice resulted in F2 progeny with 22% population exhibiting HL phenotype and 78% similar to WT mice. Hence, the HL inheritance in mice directs to a ratio of approximately 3:1. Further, HL phenotype is observed in both the male and female mice, clearly indicating that the HL phenotype follows autosomal recessive inheritance pattern (supplementary figure1).

### 3.2 Mutant mice develop cyclic alopecia

HL mice were distinguishable from their WT littermates through the entire phase of first hair cycle (postnatal days, P0 to P28). HL mice at P0 were weak with dry and wriggled skin (figure 1A). At P16, differences in coat hue was apparent and the dorsal view of mutant mice showed grey curly pelage in comparison with that of their WT littermates (figure 1A). The hair phenotype at P19 showed apparent hair loss on the head and neck regions (figure 1A). Further, the hair loss had progressively advanced in P28 and dorsal view of HL mice revealed loss of hair from head to tail region (figure 1A). HL mice were weak and have significant reduced body weights and lengths compared with that of their WT littermates (figure 1B-C) during the first phase of hair cycle formation (P0 to P28).

**Figure 1.**
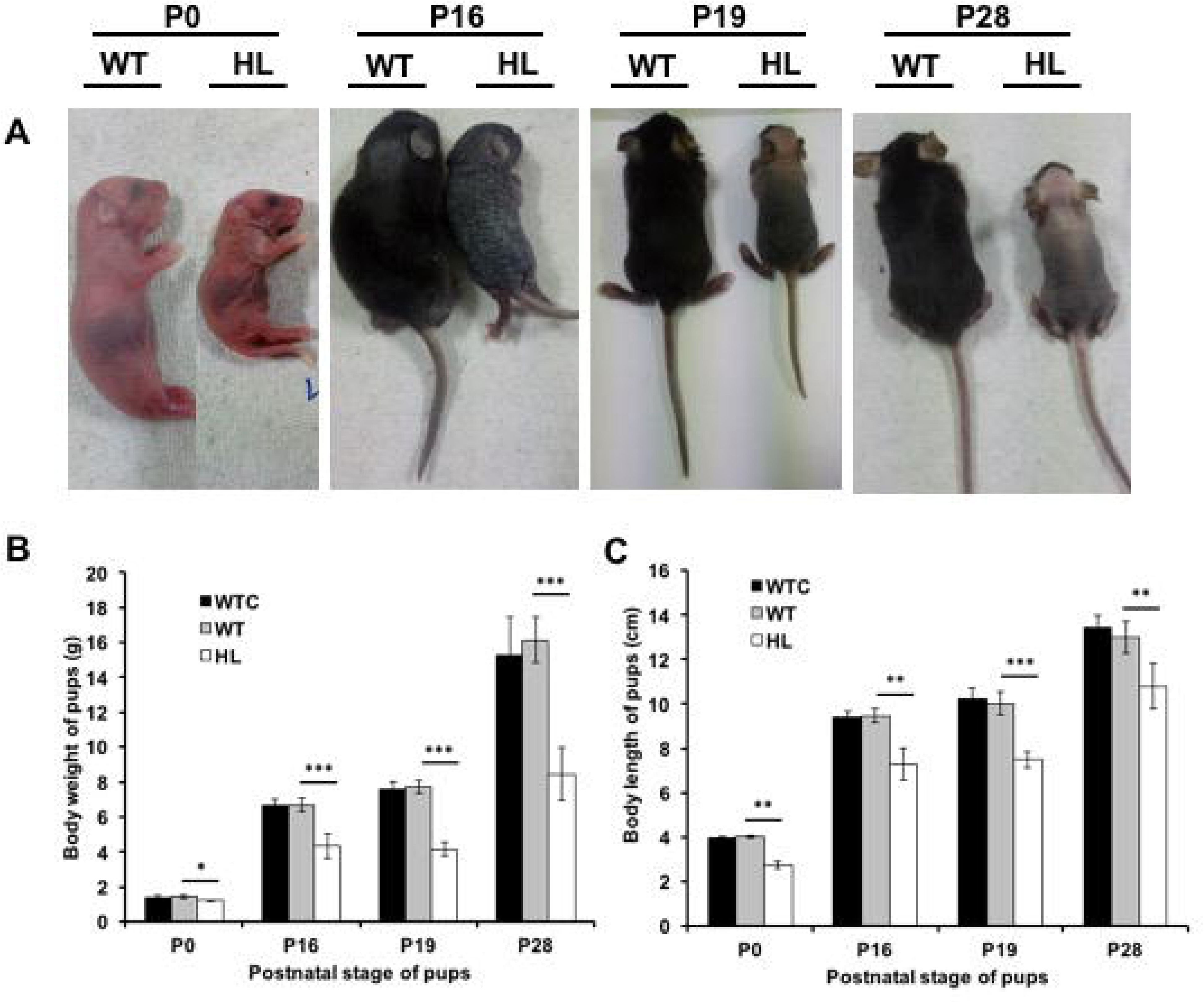
Wildtype and mutant mice at different stages of hair cycle. A) Macroscopic images of wildtype (WT) and mutant (HL) mice dorsal skin were photographed in the first hair cycle phase (P0, P16, P19 and P28) showed significant differences in the hair phenotype in HL mice in comparison with that of their WT littermates. B) Body weights were measured at postnatal days P0, P16, P19 and P28 showed significant decrease in the HL mice compared to that of their WT littermates. C) Body lengths were measured at postnatal days P0, P16, P19 and P28 showed significant decrease in the HL mice compared to that of their WT littermates. * indicates p-value less than or equal to 0.05, ** indicates p-value less than or equal to 0.001, *** indicates p-value less than or equal to 0.0001.

### 3.3 Mutant mice showed delay in the progression of catagen phase during first hair cycle formation

Hair follicle formation occurs during late embryogenesis and gets completed by postnatal day, P17 in the mice. This period of growth is known as anagen. Hair follicle undergoes regression phase known as catagen (P18-P20) where some of its lower part undergoes apoptosis. Later, the regressed hair follicle enters into resting phase called telogen (P21-P29) (Stenn and Paus 2001). Due to regenerative capability of hair-follicle stem cells, the cyclic growth of hair persists during the postnatal life (Fuchs *et al*. 2001). To investigate the hair development, histo-morphological analysis was performed for the WT and HL mice skin sections at postnatal days P0, P16, P19 and P28. At P0 and P16, the HL sections were morphologically normal in comparison with that of their WT littermates, indicating a completion of anagen phase of hair follicle morphogenesis (figure 2). Abnormal hair follicles were clearly spotted in the HL dorsal skin sections by postnatal day, P19 (figure 2). WT hair follicles are in catagen phase while the HL follicles showed delayed catagen entry with large hair bulb structures. At P28, WT hair follicles showed entry into new anagen phase but in the HL skin sections, extension of the downward growth of follicles were observed (figure 2). Taken together, morphological and macroscopic observations suggest that mutant mice show abnormal development in the first hair cycle formation which led to hairloss phenotype.

**Figure 2.**
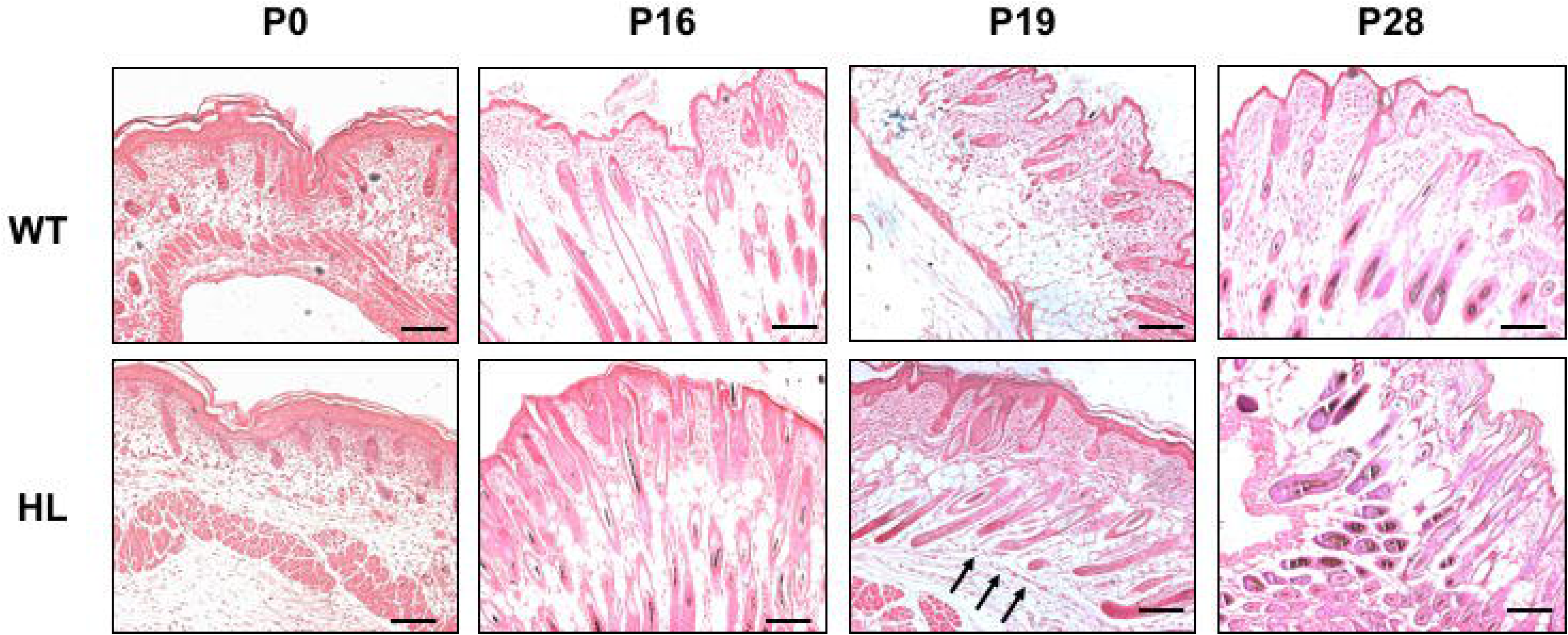
Histo-morphological examination of skin sections in mutant and wildtype mice. Histological dorsal skin sections of wildtype (WT) and mutant (HL) mice at different indicated postnatal days (P0, P16, P19 and P28). Black arrows indicate hair follicles showing delay in catagen entry. Arrows point to hair follicles exhibiting delayed regression. Magnification, 100X and scale bar represents 100 μm.

### 3.4 RNA-sequencing analysis classified stage-wise expression of genes that are associated with hair cycle formation

RNA sequencing was performed on skin tissue samples that were taken from mutant (HL) mice and it’s corresponding wildtype (WT) littermates to find the transcripts of interest that correspond to different phases of the first hair growth cycle represented by postnatal days P0, P16, P19 and P28. Gene Ontology analysis was performed for the significant differentially expressed genes (DEGs) in HL mice in comparison to that of their WT littermates (DEGs that expressed significantly at all postnatal days were considered). Using DAVID platform, pathway and gene ontology enrichment analysis was implemented for generating the differentially regulated genes. Common upregulated and down regulated genes that were expressed in all the stages of hair follicle formation were represented in table 1 and table 2. Resultant biological processes that were affected belonged to keratnization and epithelial cell differentiation processes (figure 3A). Also, cellular component analysis showed majority of genes belonged to keratin filaments and intermediate filaments (figure 3B). String analysis showed important network of interactions between the protein where most of these significant genes expressed (common among all stages of first hair cycle formation) showed plausible interactions among their protein counterparts (figure 3C). Significant DEGs that were involved in the keratinization, keratinocyte differentiation, epidermis and skin development (figure 4A) were analyzed for their role in the biological process ‘hair follicle development’ and were considered for further validation using quantitative PCR (figure 4B). We have focused and validated genes that are particularly important to hair follicle formation during first phase of hair cycle. Msx2 expression has been observed during the anagen and catagen stages of hair follicle formation and is essential for the hair shaft differentiation (Ma *et al*. 2003). Downstream effector molecule of Msx2, Lef1, expressed during anagen plays an essential role in terminal differentiation during hair follicle formation (Huelsken *et al*. 2001). Msx2 mRNA levels were significantly down regulated in HL mice in comparison to that of their WT littermates (figure 4C). LEF1 mRNA levels were downregulated in a few mice (n=4) but when all the mice were considered (n=6), there was no change in the expression levels between HL and their WT counterparts (figure 4C). Alox15 expression is found in the keratinocytes and dermal adipocytes of dorsal skin of mice (Kim *et al*. 2018). In hair loss mutant mice, Alox15 mRNA levels were significantly up regulated in late anagen (P16) and telogen (P28) stages (figure 4C). Previous study has shown that overexpression of Krt16 disrupts the keratin filaments formation in keratinocytes of skin affecting the epidermis in mice (Coulombe *et al*. 1995). QPCR analysis showed significant upregulation of Krt16 in all the stages of first hair cycle formation P0, P16, P19 and P28 (figure 4C) in the mutant mice. Krt6b is expressed in the outer root sheath of the hair follicles and glandular epithelium (Tyner *et al*. 1985). Significant up regulation of Krt6b mRNA levels at all stages (anagen, catagen and telogen) in HL mice in comparison to that of their wildtype counterparts was found (figure 4C). Fetub is expressed in the skin of mice and the function of the gene during follicle formation is not known yet (Li *et al*. 2017). Quantification of Fetub mRNA levels showed up regulation in the mutant mice at all stages (figure 4C).

**Figure 3.**
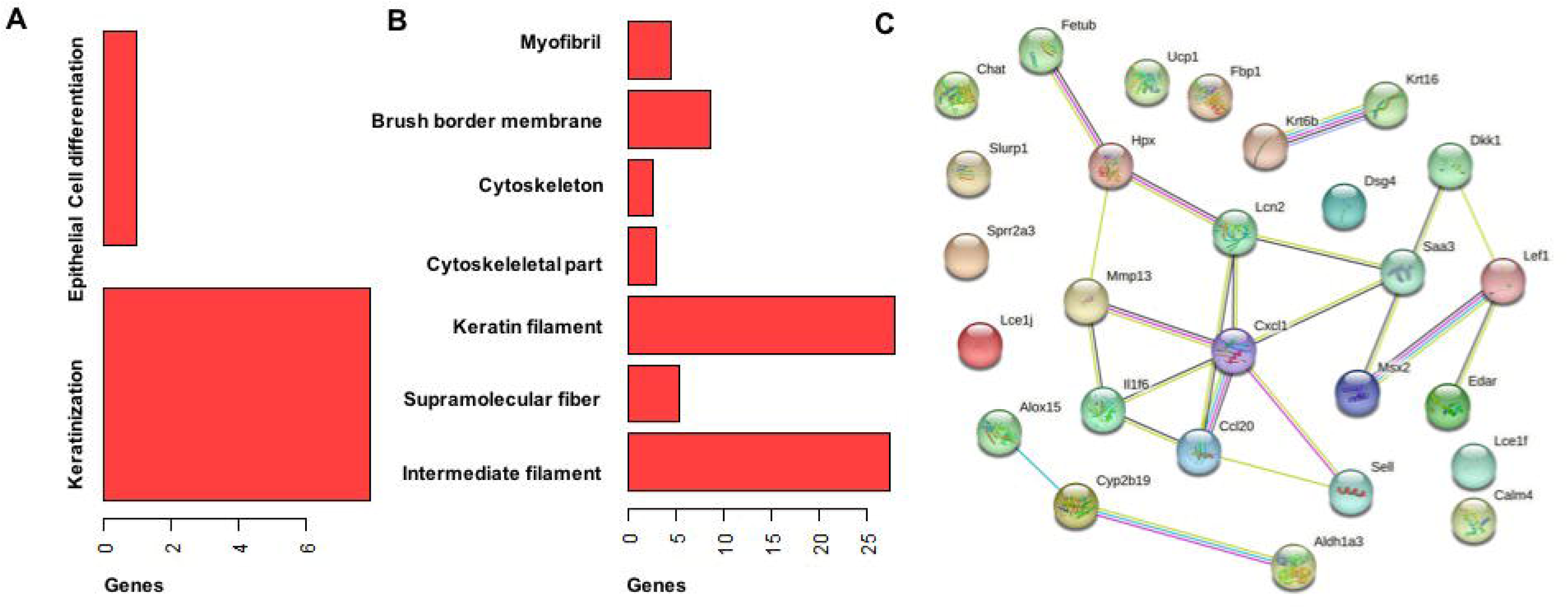
Pathway and Gene Ontology Enrichment analysis for differentially regulated genes using DAVID. Significant statistical scores from differential regulated genes and further Gene enrichment analysis indicated significant (A) Biological process and (B) Cellular component analysis. (C) Protein interacting network analysis by STRING for the most significant genes expressed (common among all stages of first hair cycle formation) showed plausible interactions among these genes.

**Figure 4.**
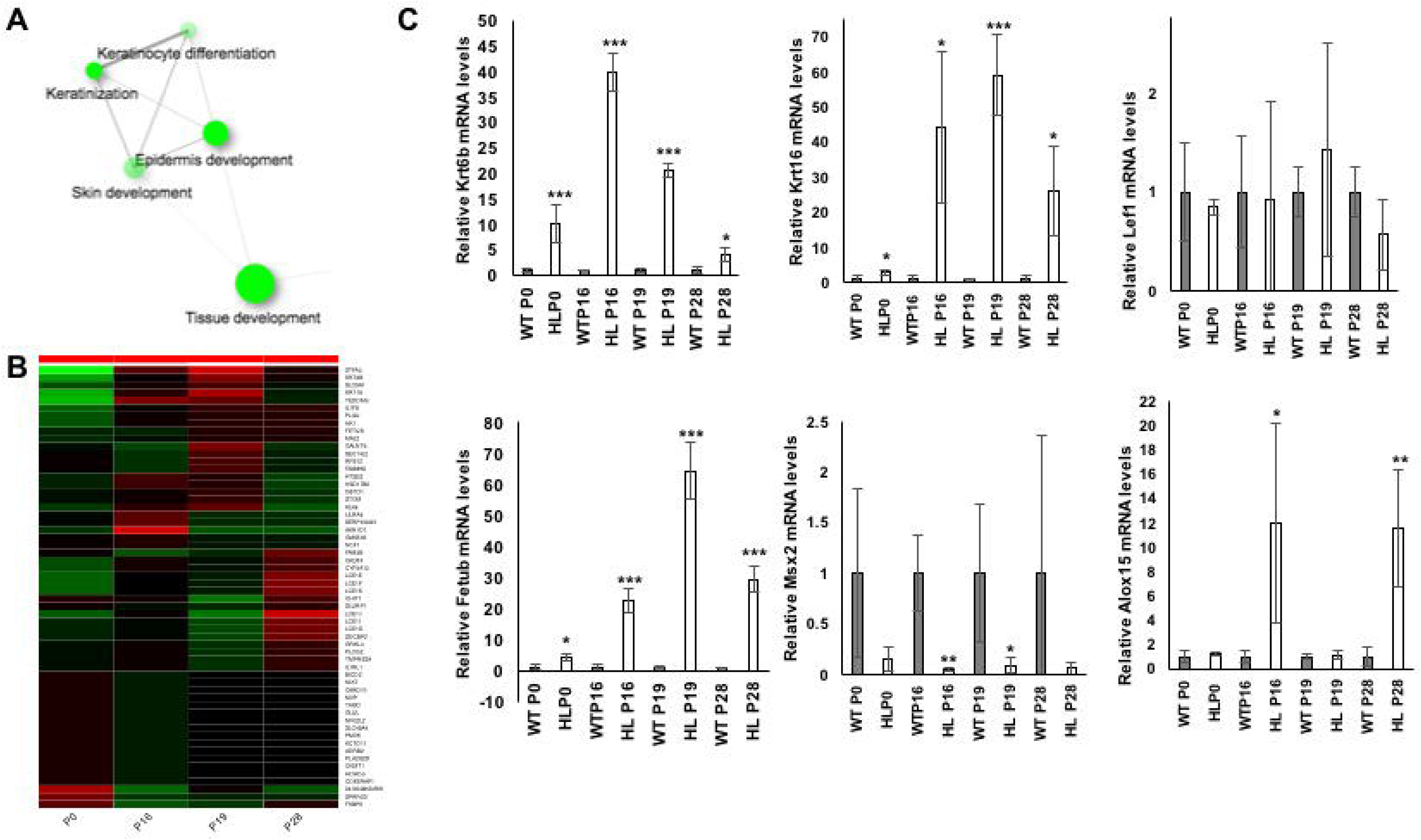
RNA sequencing and analysis for the identification of hair loss associated genes at different stages of first hair cycle. A) Significant biological process involved for the differentially regulated genes at the first hair cycle phase (P0, P16, P19 and P28). B) Heat map of significant differentially regulated genes associated with hair follicle formation stage wise (anagen (P0, P16), catagen (P19) and telogen (P28)). C) Quantitative-PCR (QPCR) results of gene expression for postnatal days-P0, P16, P19 and P28 covering the first phase of hair cycle was performed. Relative mRNA levels for significant genes (Krt6b, Krt16, Lef1, Fetub, Msx2 and Alox15) and standard deviations were calculated by using replicates (n=6). RPL13 was considered as internal control. * indicates p-value less than or equal to 0.05, ** indicates p-value less than or equal to 0.001, *** indicates p-value less than or equal to 0.0001.

**Table 1.**
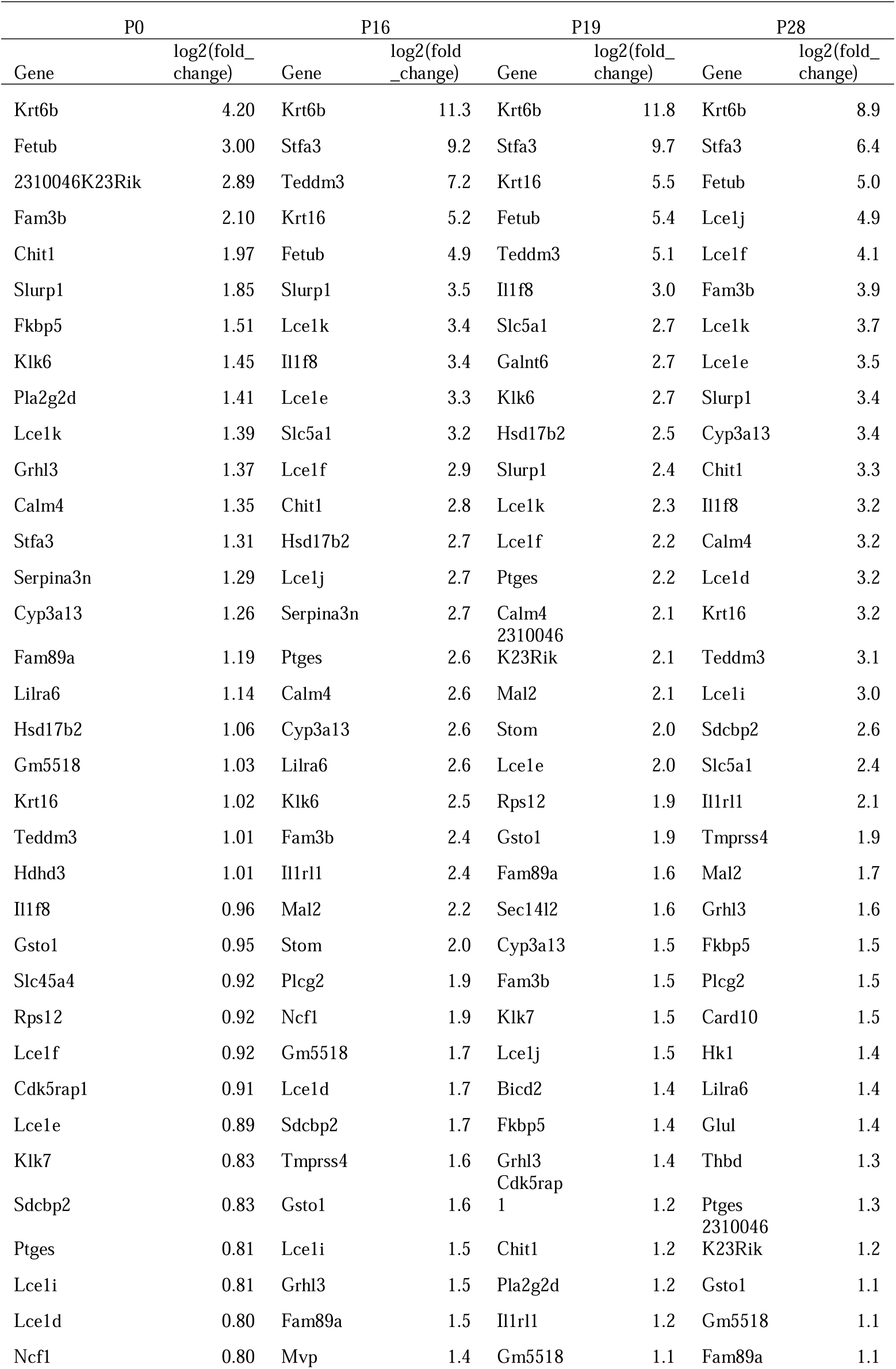
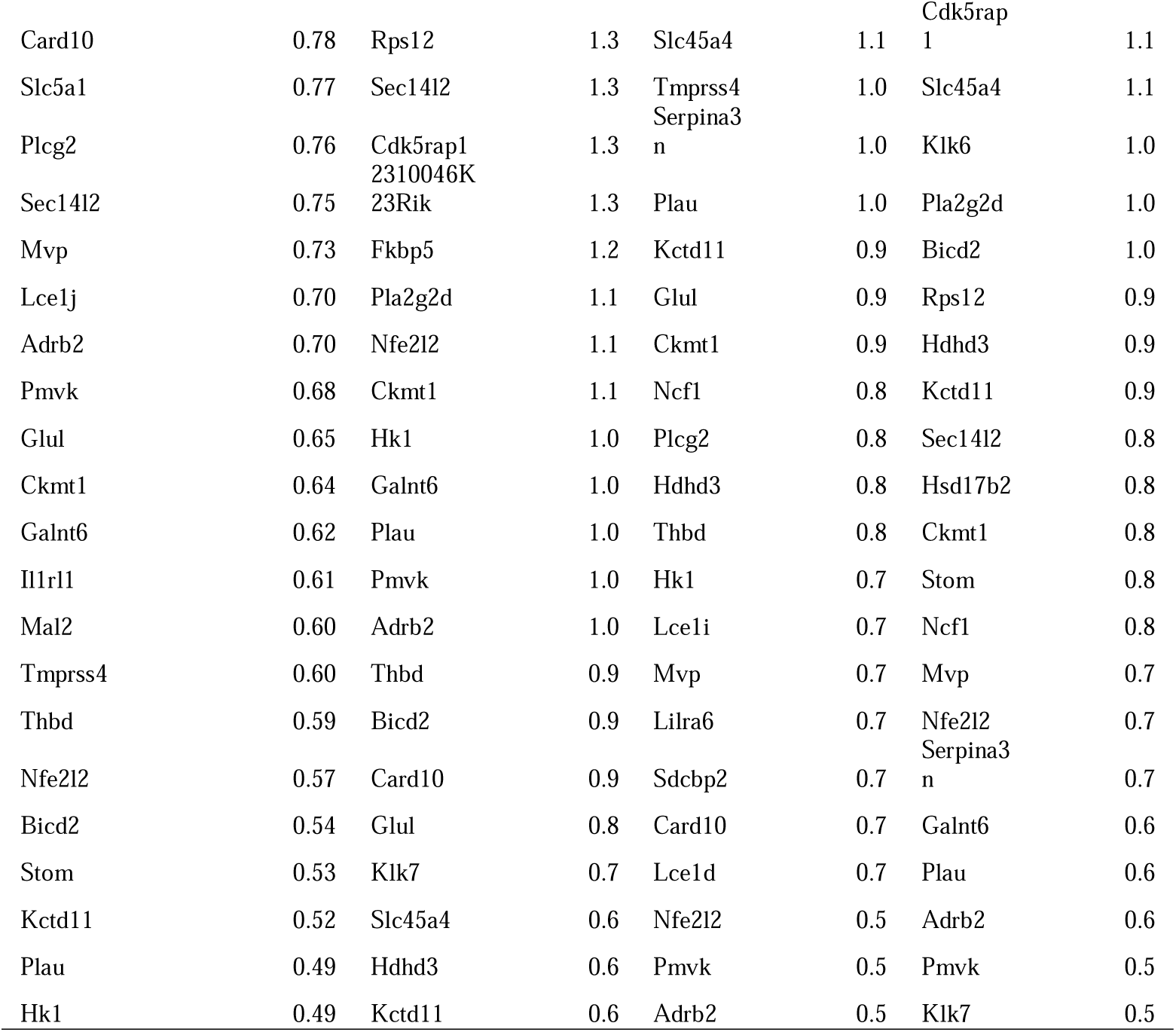
A comparative list of common upregulated genes with log2FC values in mutant mice in comparison to the wildtype at different stages of first hair cycle formation (P0, P16, P19 and P28).

**Table 2.** A comparative list of common downregulated genes with log2FC values in mutant mice in comparison to the wildtype at different stages of first hair cycle formation (P0, P16, P19 and P28).

| P0 |  | P16 |  | P19 |  | P28 |  |
| --- | --- | --- | --- | --- | --- | --- | --- |
| Gene | log2(fold<br>_change) | Gene | log2(fold<br>_change) | Gene | log2(fold<br>_change) | Gene | log2(fold<br>_change) |
| Ndufa7 | -0.5 | Cox8a | -0.5 | Cox8a | -0.5 | Nenf | -0.8 |
| Nenf | -0.6 | Ndufa7 | -0.5 | Pdcd5 | -0.6 | Lce1m | -0.8 |
| Smpd1 | -0.7 | Nenf | -0.6 | Ndufa7 | -0.9 | Ndufa7 | -0.9 |
| Cox8a | -0.8 | Cox5b | -0.8 | Nenf | -0.9 | Tmem229a | -1.0 |
| Lce1m | -0.8 | Ndufs6 | -1.1 | Muc15 | -0.9 | Smpd1 | -1.0 |
| Atp5e | -0.8 | Atp5e | -1.2 | Ndufs6 | -0.9 | Ndufs6 | -1.1 |
| Cox5b | -0.8 | Muc15 | -1.2 | Cox5b | -1.1 | Gm22009 | -1.1 |
| Mlana | -0.8 | Smpd1 | -1.5 | Hist1h4<br>c | -1.1 | Atp5e | -1.2 |
| Scarna2 | -0.8 | Btc | -1.5 | Slc6a19 | -1.2 | Muc15 | -1.3 |
| Pdcd5 | -0.9 | Hist1h4<br>c | -1.5 | Scarna2 | -1.2 | Btc | -1.4 |
| Gm22009 | -0.9 | Pdcd5 | -1.5 | Atp5e | -1.2 | Cox8a | -1.4 |
| Ndufs6 | -1.0 | Tmem2<br>29a | -1.7 | Hist1h4<br>h | -1.2 | Cyp2b19 | -1.5 |
| Muc15 | -1.0 | Hist1h4 | -1.7 | Gm220 | -1.3 | Cox5b | -1.5 |

|  |  | h |  | 09 |  |  |  |
| --- | --- | --- | --- | --- | --- | --- | --- |
|  |  | Gm220 |  |  |  |  |  |
| Hist1h4c | -1.0 | 09 | -2.0 | Smpd1 | -1.4 | Skint10 | -1.7 |
|  |  |  |  | Hist3h2 |  |  |  |
| Tmem229a | -1.1 | Scarna2 | -2.0 | ba | -1.4 | Scarna2 | -1.9 |
| Hist3h2ba | -1.2 | Rpph1 | -2.2 | Mlana | -2.0 | Pdcd5 | -1.9 |
| Skint10 | -1.4 | Skint10 | -2.2 | Rpph1 | -2.1 | Hist1h4c | -2.1 |
|  |  | Hist3h2 |  | Cyp2b1 |  |  |  |
| Serpina3b | -1.4 | ba | -2.5 | 9 | -2.3 | Rpph1 | -2.2 |
| Hist1h4h | -1.5 | Dapl1 | -2.5 | Skint10 | -2.8 | Serpina3b | -2.2 |
|  |  |  |  | Tmem2 |  |  |  |
| Btc | -1.6 | Mlana | -2.6 | 29a | -2.9 | Hist1h4h | -2.5 |
|  |  | Cyp2b1 |  |  |  |  |  |
| Dapl1 | -2.1 | 9 | -2.7 | Btc | -2.9 | Hist3h2ba | -2.7 |
| Slc6a19 | -2.5 | Slc6a19 | -3.1 | Dapl1 | -3.3 | Mlana | -3.1 |
| Rpph1 | -2.7 | Lce1m | -3.3 | Lce1m | -4.3 | Dapl1 | -3.3 |
|  |  | Serpina |  |  |  |  |  |
| Cyp2b19 | -3.1 | 3b | -4.5 | Zfp125 | -4.7 | Zfp125 | -5.2 |
|  |  |  |  | Serpina |  |  |  |
| Zfp125 | -4.0 | Zfp125 | -6.4 | 3b | -6.8 | Slc6a19 | -9.8 |

### 3.5 Alox15 and Krt6b are overexpressed in hairloss skin sections

QPCR validation showed significant overexpression of Alox15 and Krt6b at different stages of hair follicle formation, we assessed protein levels of the same. Immunoblot analysis showed increased expression levels of Alox15 at P16 and P28 stages in skin sections of hairloss mice (figure 5A,C). Krt6b was increased in mutant mice at all stages (P0, P16, P19 and P28) of the hair cycle formation in comparison to that of their wildtype counterparts (figure 5E,F). Msx2 protein is essential in the formation of hair follicle and its absence leads to hairloss phenotype (Ma *et al*. 2003). Immunoblot analysis revealed significant low Msx2 levels in mutant mice at P19 stage (figure 5B,D).

**Figure 5.**
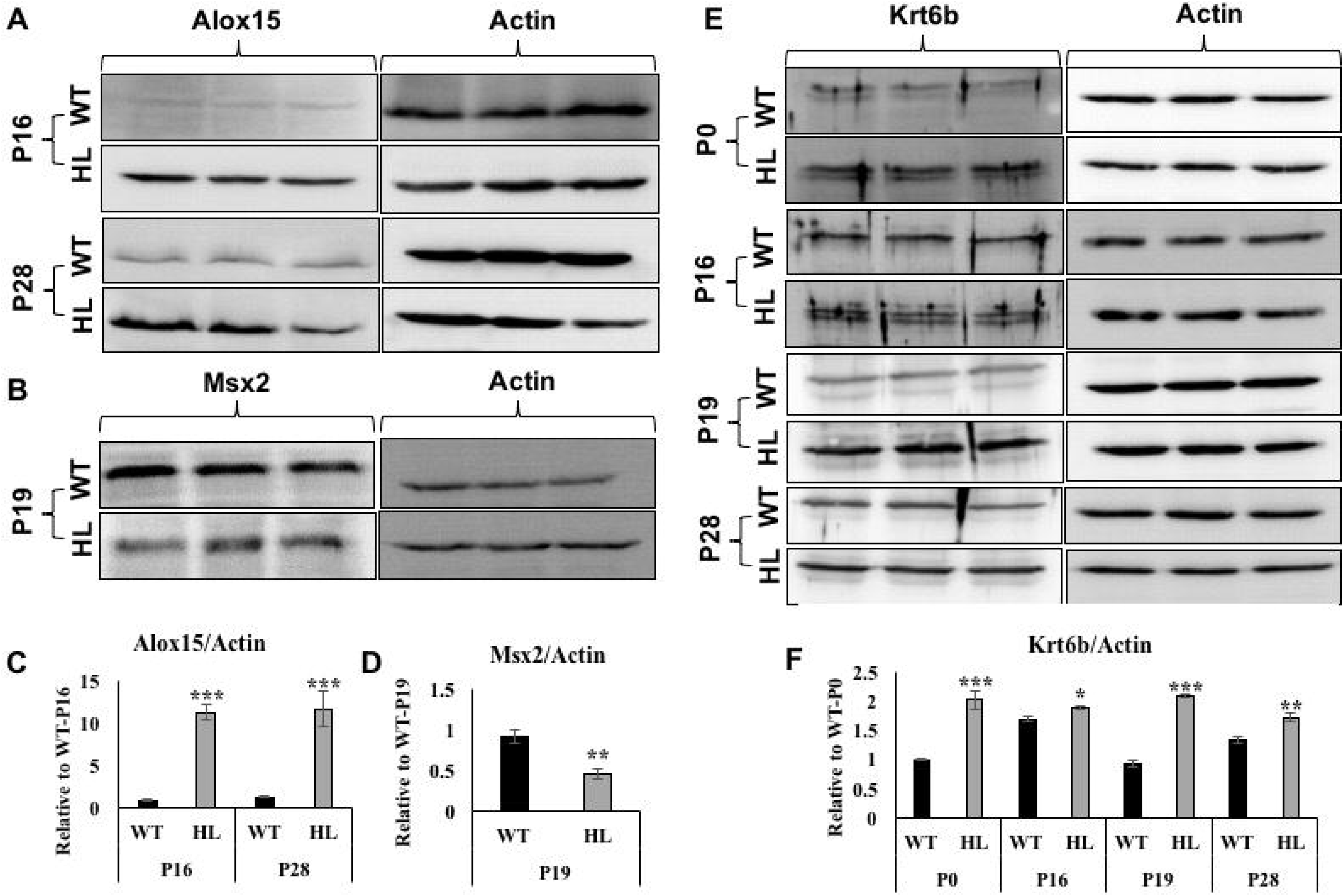
Immunoblot analysis of Alox15, Msx2 and Krt6b at different stages of hair cycle formation. Immunoblot analysis of differentially expressed significant markers-A) Alox15 B) Msx2 and E) Krt6b were analyzed in hairloss and wildtype mice. Quantification of the immunoblot and its analysis was done for C) Alox15, D) Msx2 and F) Krt6b. Data presented were mean ± SEM (n =3). * indicates p-value less than or equal to 0.05, ** indicates p-value less than or equal to 0.001, *** indicates p-value less than or equal to 0.0001.

### 3.6 Krt6b is overexpressed in epidermal region of the mutant mice

Krt6b is overexpressed in all mutant mice studied at all stages of hair cycle formation suggesting it as a novel biomarker associated with hairloss condition. Immuno-histochemical evaluation revealed expression of krt6b in hair shafts of wildtype mice whereas hair shaft of mutant mice did not show any expression at P0 stage (figure 6). The epidermal region of mutant mice showed overexpression of krt6b at P0, P16, P19 and P28 stages whereas in wildtype skin sections the expression is restricted to hair shafts (figure 6).

**Figure 6.**
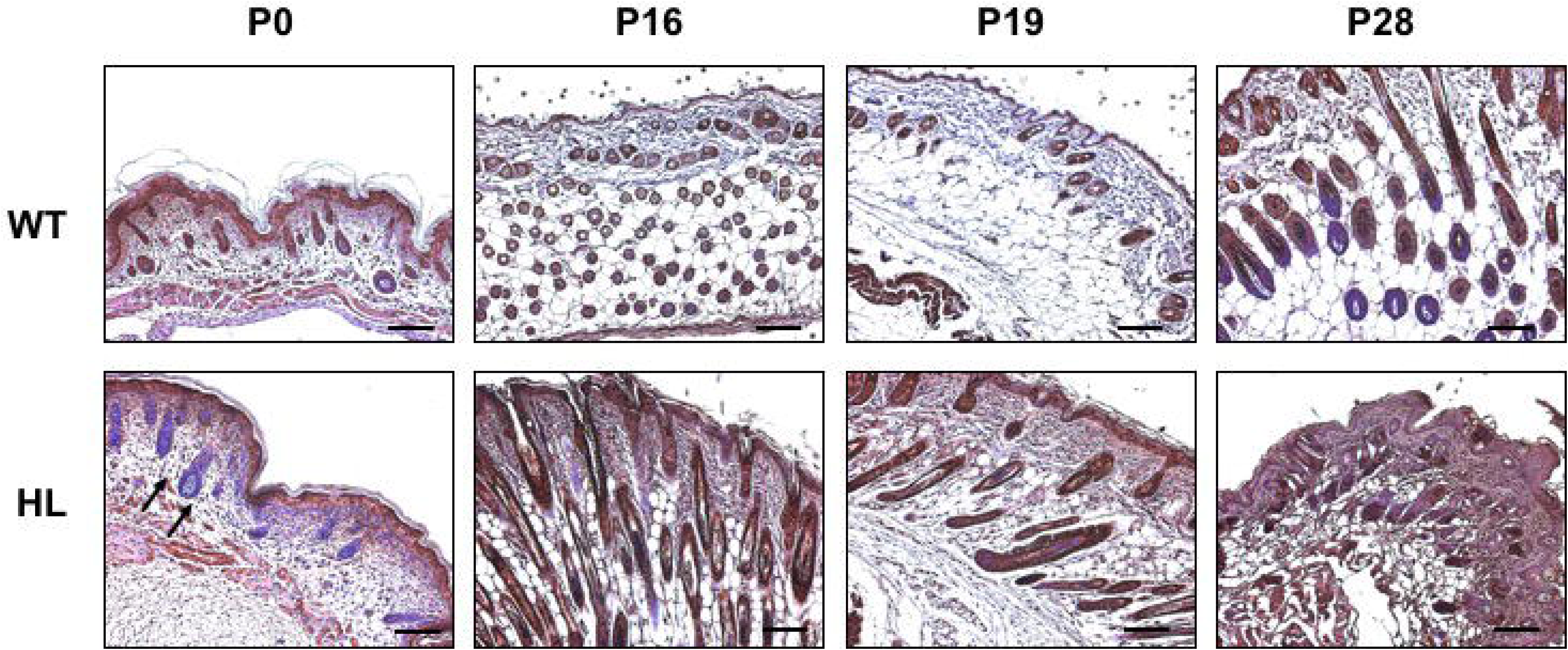
Immunohistochemistry (IHC) analysis of krt6b at different stages of first hair cycle. IHC was performed on skin sections of wildtype (WT) and mutant (HL) C57BL/6 mice at different postnatal days (P0, P16, P19 and P28). Black arrows indicate the hair shafts devoid of Krt6b staining at P0 stage in HL mice. Scalebar=50 μm, magnification-100x.

## 4. Discussion

In-house inbred mouse strain, C57BL/6, developed spontaneous mutation leading to hair loss condition that displayed pathophysiological condition of cyclic alopecia. We have analyzed the histo-morphological and molecular events in the hair loss mutant (HL) mice. These analyses revealed that there is a defect in the first cycle of hair follicle formation. Macroscopic examination of skin sections showed changes initiated at late anagen phase with grey curly pelage. Histo-morphological examination showed changes in the catagen-telogen transition with large hair bulb like structures and changes in regression of hair follicle development. The delay in the change of the phase has led to structural abnormalities during catagen phase leading to cyclic alopecia phenotype which is characterized by the delayed stages and eventually resulting in hair loss condition. It also displayed asynchronous cycling in the skin at different body regions.

Many studies have shown the occurrence of cyclic alopecia in several genetic conditions of mice. Msx2 is one of the homeodomain transcriptional factors that controls development of the hair follicles, heart, teeth, skull and brain (Zhao *et al*. 2012). Msx2 knockout mice did not show any other phenotype except for the hair loss. *Msx2^-/-^*mice showed hair loss phenotype due to hair cycle defect. Loss of Msx2 leads to abnormal hair shaft differentiation and its breakage attributing to hair loss phenotype (Ma *et al*. 2003). Our results of low expression levels of MSX2 in HL mice are in agreement with the previous study of *Msx2^-/-^*mice where hair loss phenotype was observed due to hair cycle defect. Alox15 is an enzyme involved in the generation of bioactive lipid metabolites (Singh and Rao 2019). Alox15 is abundantly expressed in keratinocytes and plays an important role in skin homeostasis. Alox15 knockout mice showed hair loss at the age of 16 weeks with increased pro-inflammatory and necroptic response in the dermal adipocyte region (Kim *et al*. 2018). Contrary to these observations, our data showed increase in the Alox15 mRNA and protein levels in the HL mice. One of the study reported that Alox15 levels get increased in the patients suffering from critical illness with low body weights (Goossens *et al*. 2017). In our present study, HL mice had lower body weights and may be Alox15 levels were increased to promote lipid accumulation which has to be further validated. Genetic studies of several forms of alopecia showed novel loci/genes that control the hair growth and formation. Mutations in genes – Corneodesmosin (CDSN), hairless (HR) and desmoglein4 (DSG4) results in hairloss condition which is known to be autosomal recessive inheritance (Kljuic *et al*. 2003; Levy-Nissenbaum *et al*. 2003). Although there are number of studies suggesting that the mutations in genes lead to hair loss condition(Kljuic *et al*. 2003; Levy-Nissenbaum *et al*. 2003), present study results suggest that there might be involvement of combination of more number of mutated genes leading to hair loss condition which has to be further explored.

Keratin gene mutations and formation of aggregates are known to be associated with alopecia condition (Chamcheu *et al*. 2011). In comparison to the wildtype, mutant mice showed upregulation of krt6b and krt16 levels at all stages of first hair cycle. Further, the epidermal and dermal counterparts showed overexpression of krt6b in mutant mice suggesting the possibility of its association in aggregation and abnormality in keratinocyte formation and differentiation which needs to be studied in detail in future. Most of the genes associated with keratin filaments that play an important role in keratinization have also been upregulated in the current study. Future strategies involving therapies that target these upregulated keratin proteins such as use of retinoic acid (Li and Törmä 2013) to control the expression of keratin proteins can be studied and further can be implemented in therapeutic use.

Taken together, the present study proposes a model for relative hair follicle cycling analysis. Abnormal hair follicle cycling and morphology in the HL mice affected the hair shaft growth and development due to delay in the progression through the first cycle of hair follicle process. The hair loss mouse model in the present study may play an important role in the identification, evaluation and treatment strategies for alopecia condition. Also, further studies are necessary to investigate the role of inflammation associated with the hair loss condition in these mice.

## Supporting information

Full length blots

## Acknowledgements

The authors thank Jerald Mahesh Kumar and Jyothi Lakshmi for technical help in the animal breeding studies. This study was supported by the grant from the Department of Biotechnology (BT/PR15320/BRB/10/1453/2015), Govt. of India.

## Conflict of interest

The authors declare no competing financial interest.

## Notes

https://www.ncbi.nlm.nih.gov/geo/query/acc.cgi?acc=GSE220769

### Competing Interest Statement

The authors have declared no competing interest.

## References

1. Cai J, Lee J, Kopan R and Ma L 2009 Genetic interplays between Msx2 and Foxn1 are required for Notch1 expression and hair shaft differentiation. Dev Biol 326 420–430

2. Chamcheu JC, Siddiqui IA, Syed DN, Adhami VM, Liovic M et al. 2011 Keratin gene mutations in disorders of human skin and its appendages. Arch Biochem Biophys 508 123–137

3. Cheng JB, Sedgewick AJ, Finnegan AI, Harirchian P, Lee J et al. 2018 Transcriptional Programming of Normal and Inflamed Human Epidermis at Single-Cell Resolution. Cell Rep 25 871–883

4. Coulombe PA, Bravo NS, Paladini RD, Nguyen D and Takahashi K 1995 Overexpression of human keratin 16 produces a distinct skin phenotype in transgenic mouse skin. Biochem Cell Biol 73 611–618

5. Eckhart L, Lippens S, Tschachler E and Declercq W 2013 Cell death by cornification. Biochim Biophys Acta 1833 3471–3480

6. Fuchs E, Merrill BJ, Jamora C and DasGupta R 2001 At the roots of a never-ending cycle. Dev Cell 1 13–25

7. Goossens C, Vander Perre S, Van den Berghe G and Langouche L 2017 Proliferation and differentiation of adipose tissue in prolonged lean and obese critically ill patients. Intensive Care Med Exp 5 16

8. Huelsken J, Vogel R, Erdmann B, Cotsarelis G and Birchmeier W 2001 beta-Catenin controls hair follicle morphogenesis and stem cell differentiation in the skin. Cell 105 533–545

9. Joost S, Zeisel A, Jacob T, Sun X, La Manno G et al. 2016 Single-Cell Transcriptomics Reveals that Differentiation and Spatial Signatures Shape Epidermal and Hair Follicle Heterogeneity. Cell Syst 3 221–237.e229

10. Kim SN, Akindehin S, Kwon HJ, Son YH, Saha A et al. 2018 Anti-inflammatory role of 15-lipoxygenase contributes to the maintenance of skin integrity in mice. Sci Rep 8 8856

11. Kljuic A, Bazzi H, Sundberg JP, Martinez-Mir A, O’Shaughnessy R et al. 2003 Desmoglein 4 in hair follicle differentiation and epidermal adhesion: evidence from inherited hypotrichosis and acquired pemphigus vulgaris. Cell 113 249–260

12. Kobielak K, Stokes N, de la Cruz J, Polak L and Fuchs E 2007 Loss of a quiescent niche but not follicle stem cells in the absence of bone morphogenetic protein signaling. Proc Natl Acad Sci U S A 104 10063–10068

13. Langbein L, Rogers MA, Winter H, Praetzel S and Schweizer J 2001 The catalog of human hair keratins. II. Expression of the six type II members in the hair follicle and the combined catalog of human type I and II keratins. J Biol Chem 276 35123–35132

14. Levy-Nissenbaum E, Betz RC, Frydman M, Simon M, Lahat H et al. 2003 Hypotrichosis simplex of the scalp is associated with nonsense mutations in CDSN encoding corneodesmosin. Nat Genet 34 151–153

15. Li C, Gao C, Fu Q, Su B and Chen J 2017 Identification and expression analysis of fetuin B (FETUB) in turbot (Scophthalmus maximus L.) mucosal barriers following bacterial challenge. Fish Shellfish Immunol 68 386–394

16. Li H and Törmä H 2013 Retinoids reduce formation of keratin aggregates in heat-stressed immortalized keratinocytes from an epidermolytic ichthyosis patient with a KRT10 mutation*. Acta Derm Venereol 93 44–49

17. Lim X and Nusse R 2013 Wnt signaling in skin development, homeostasis, and disease. Cold Spring Harb Perspect Biol 5

18. Ma L, Liu J, Wu T, Plikus M, Jiang TX et al. 2003 ’Cyclic alopecia’ in Msx2 mutants: defects in hair cycling and hair shaft differentiation. Development 130 379–389

19. Oka A, Takagi A, Komiyama E, Yoshihara N, Mano S et al. 2020 Alopecia areata susceptibility variant in MHC region impacts expressions of genes contributing to hair keratinization and is involved in hair loss. EBioMedicine 57 102810

20. Schneider MR, Schmidt-Ullrich R and Paus R 2009 The hair follicle as a dynamic miniorgan. Curr Biol 19 R132–142

21. Schweizer J, Langbein L, Rogers MA and Winter H 2007 Hair follicle-specific keratins and their diseases. Exp Cell Res 313 2010–2020

22. Singh NK and Rao GN 2019 Emerging role of 12/15-Lipoxygenase (ALOX15) in human pathologies. Prog Lipid Res 73 28–45

23. Singh S, Pavuluri S, Jyothi Lakshmi B, Biswa BB, Venkatachalam B et al. 2020 Molecular characterization of Wdr13 knockout female mice uteri: a model for human endometrial hyperplasia. Sci Rep 10 14621

24. Stenn KS and Paus R 2001 Controls of hair follicle cycling. Physiol Rev 81 449–494

25. Tyner AL, Eichman MJ and Fuchs E 1985 The sequence of a type II keratin gene expressed in human skin: conservation of structure among all intermediate filament genes. Proc Natl Acad Sci U S A 82 4683–4687

26. Yang H, Adam RC, Ge Y, Hua ZL and Fuchs E 2017 Epithelial-Mesenchymal Micro-niches Govern Stem Cell Lineage Choices. Cell 169 483–496.e413

27. Zhang B, Ma S, Rachmin I, He M, Baral P et al. 2020 Hyperactivation of sympathetic nerves drives depletion of melanocyte stem cells. Nature 577 676–681

28. Zhao J, Kawai K, Wang H, Wu D, Wang M et al. 2012 Loss of Msx2 function down-regulates the FoxE3 expression and results in anterior segment dysgenesis resembling Peters anomaly. Am J Pathol 180 2230–2239

