## Supplementary Table 1, Figure1 for "Differential expression of keratin and keratin associated proteins are linked with hair loss condition in spontaneously mutated inbred mice"

Supplementary Table 1. List of primers used for quantitative PCR

| **Gene** | **Primer Sequence 5' --> 3'** |
| --- | --- |
| Rpl13a | Fwd-TTGCTTACCTGGGGCGTCT |
|  | Rev-CCTTTTCCTTCCGTTTCTCCTC |
| Krt6b | Fwd-TTGCCACCTACAGGAAGCTG |
|  | Rev-CCCAGGCCTAAGCTGCTG |
| Krt16 | Fwd-GTCTGCTGGATGGCGAGAAT |
|  | Rev-CTCTGGCTGAAGCTGGTTGA |
| Fetub | Fwd-ACCCGTAGAAAAGTCTGTCACT |
|  | Rev-CCTCTTGGTGCAGTTACGGA |
| Msx2 | Fwd-CCACATCCCAGCTTCTAGCC |
|  | Rev-TTCCGCCTCTTGCAGTCTTT |
| Lef1 | Fwd-CGGGAAGAGCAGGCCAAATA |
|  | Rev-AGTCGACTCCTGTAGCTTCTCT |
| Alox15 | Fwd-ACGATGGAGAAGCTGATGGC |
|  | Rev-TTCTTCAGCACAGCTTTGGC |

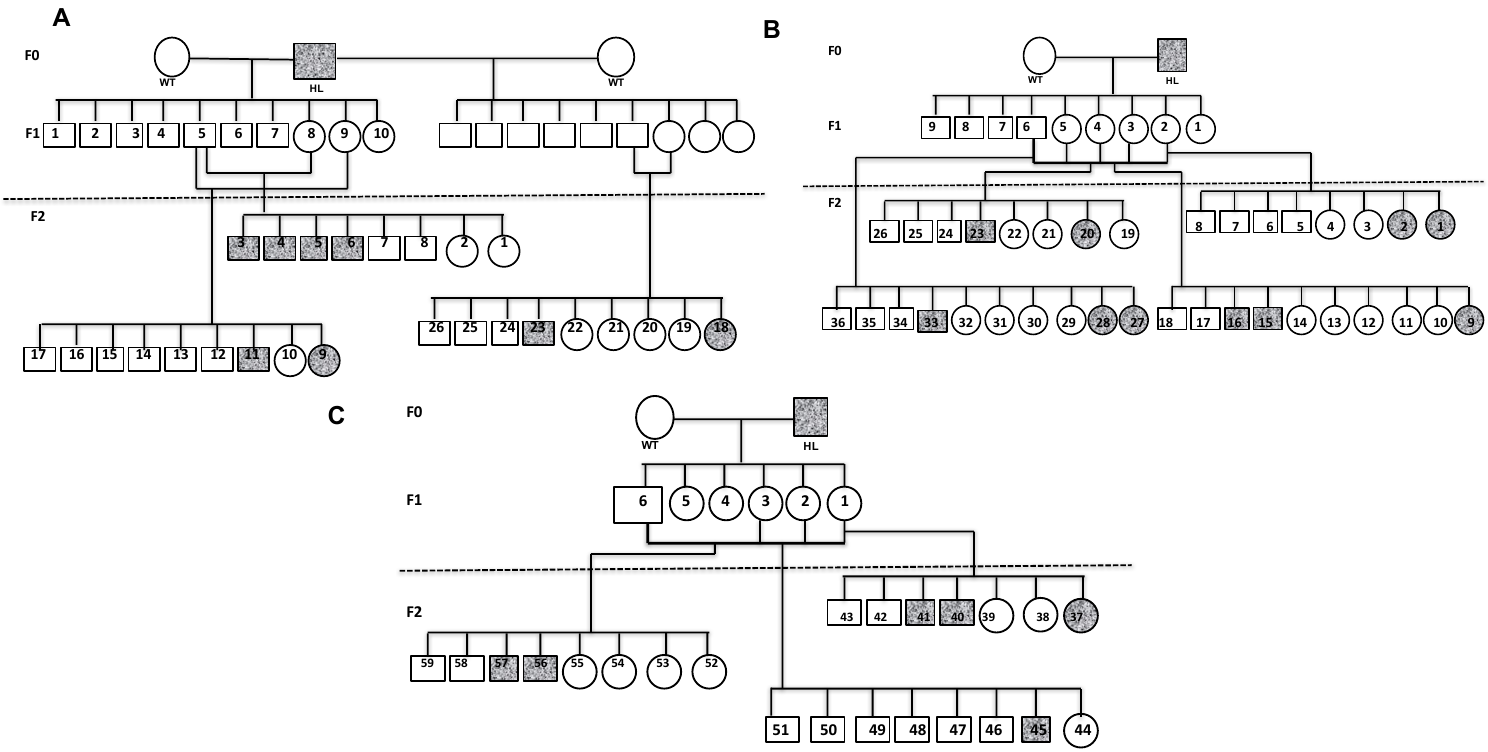

Supplementary figure 1. Hairloss phenotype in C57BL6J mice displays autosomal recessive inheritance. (A-C) Wildtype mice were mated with mutant mice (HL). HL mice were represented in grey filled ones. The resultant mice in F1 generation were heterozygous. Further littermates were crossed in F1 and nearly quarter of the progeny were found to be wild type and mutant mice.
