## Supplementary material for "Differential expression of keratin and keratin associated proteins are linked with hair loss condition in spontaneously mutated inbred mice": Full length blots

Figure 5A –P16

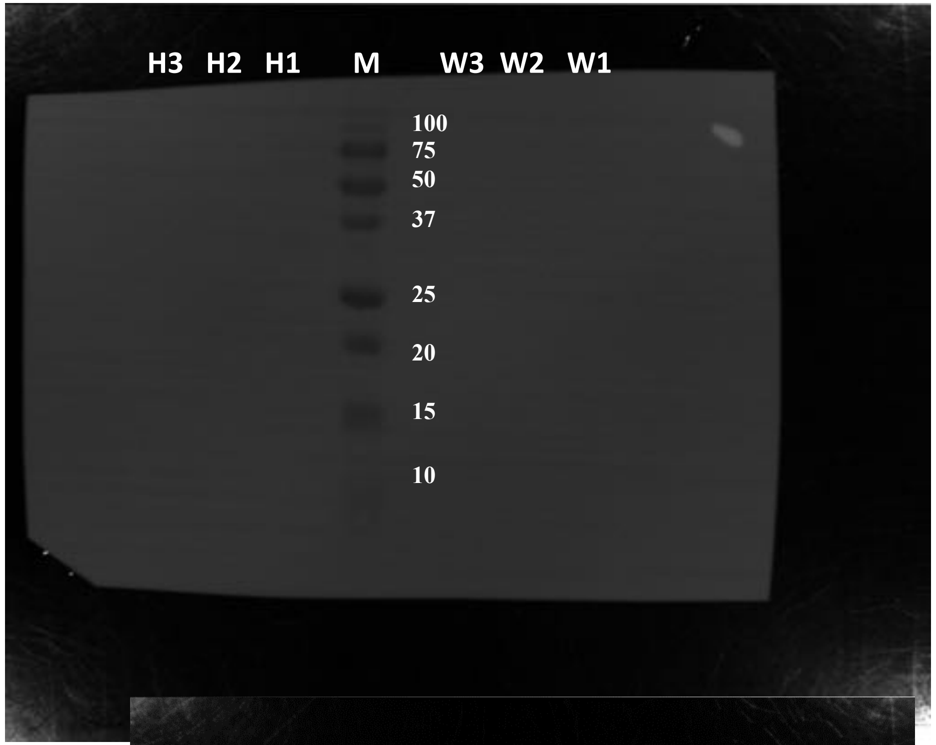


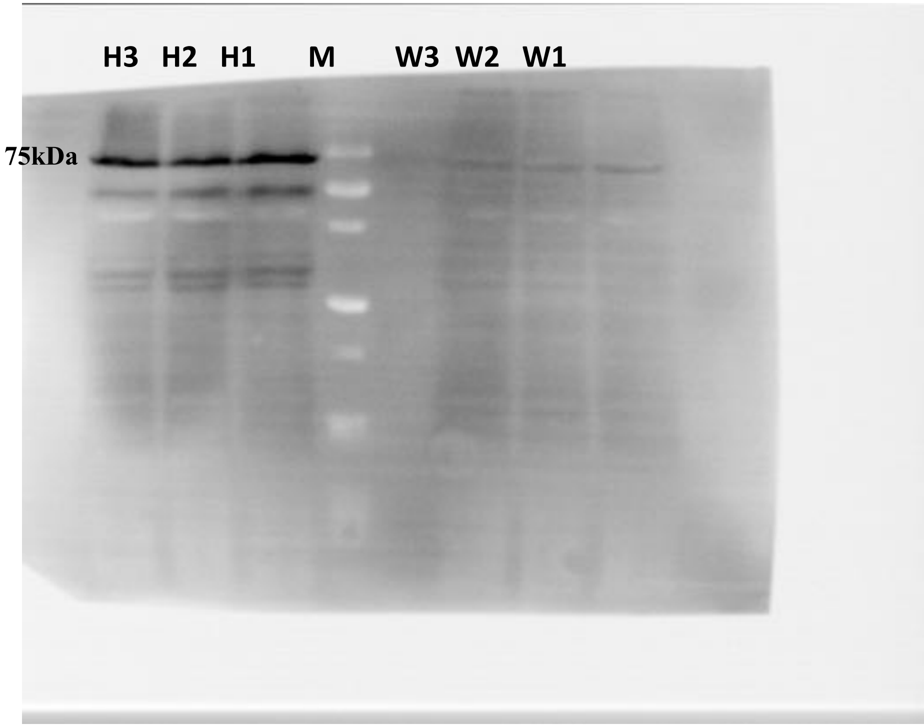


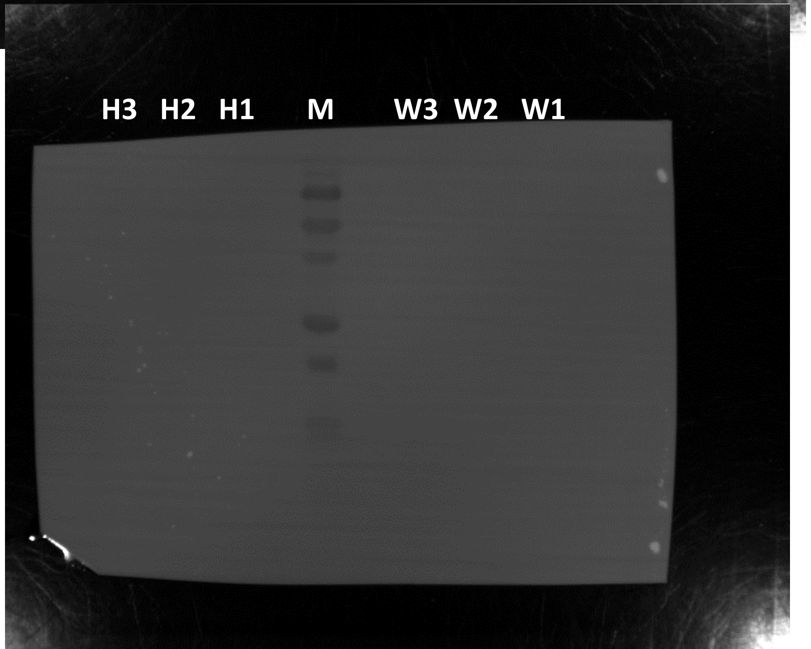


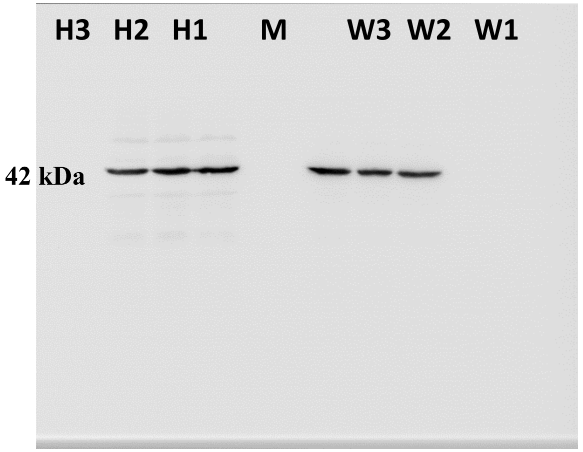


**All Blots has Marker- Biorad All Blue Prestained Protein Standard, Cat #1610373**

### Figure 5A –P28

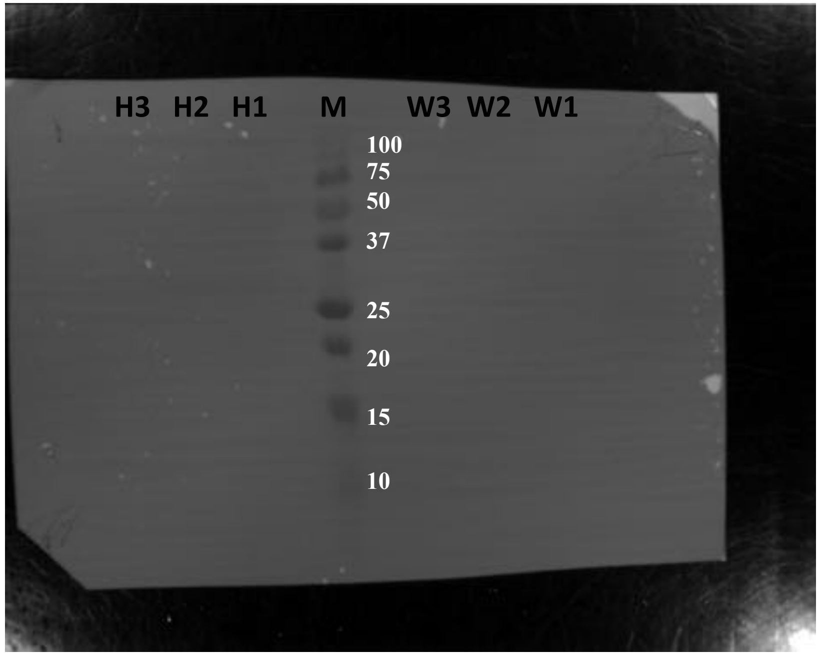


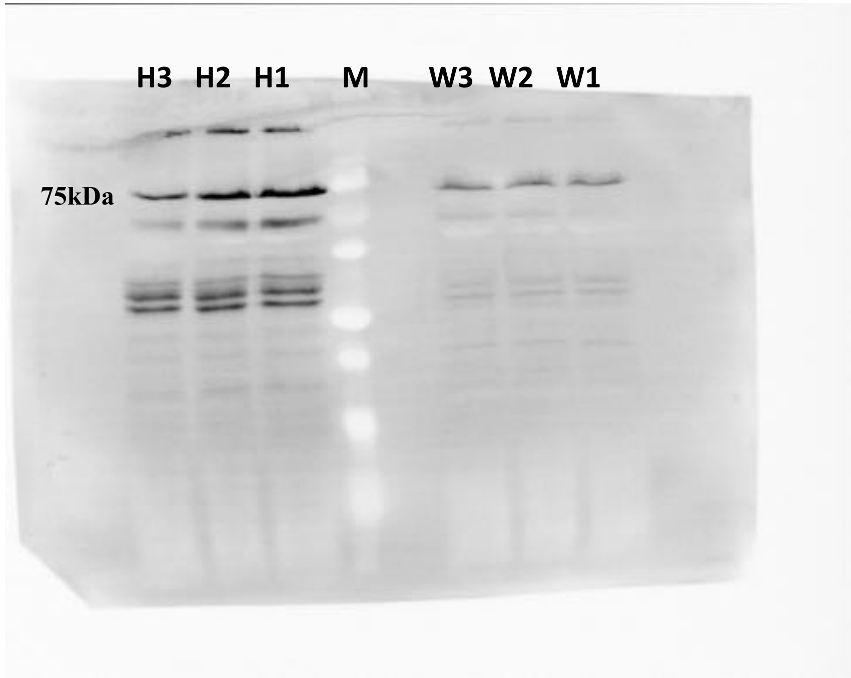


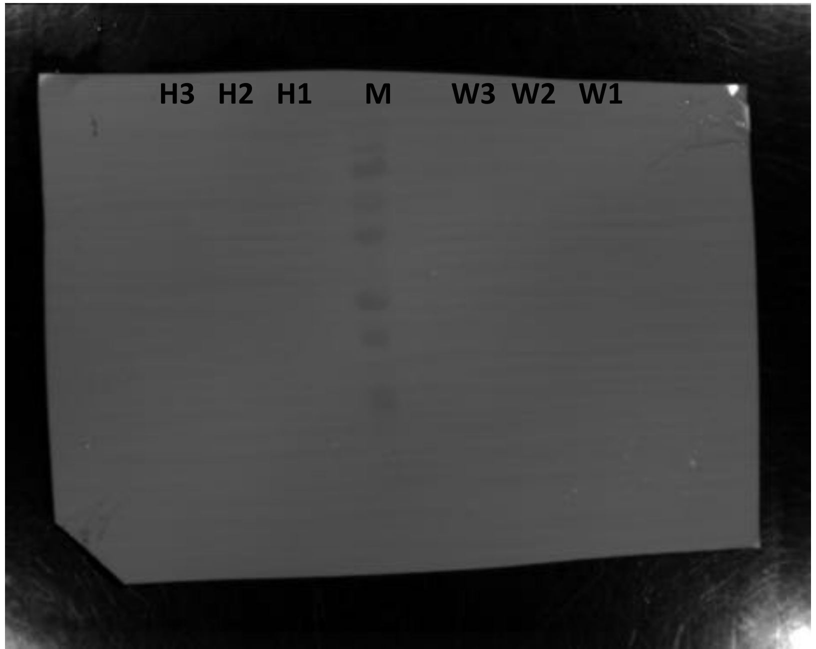


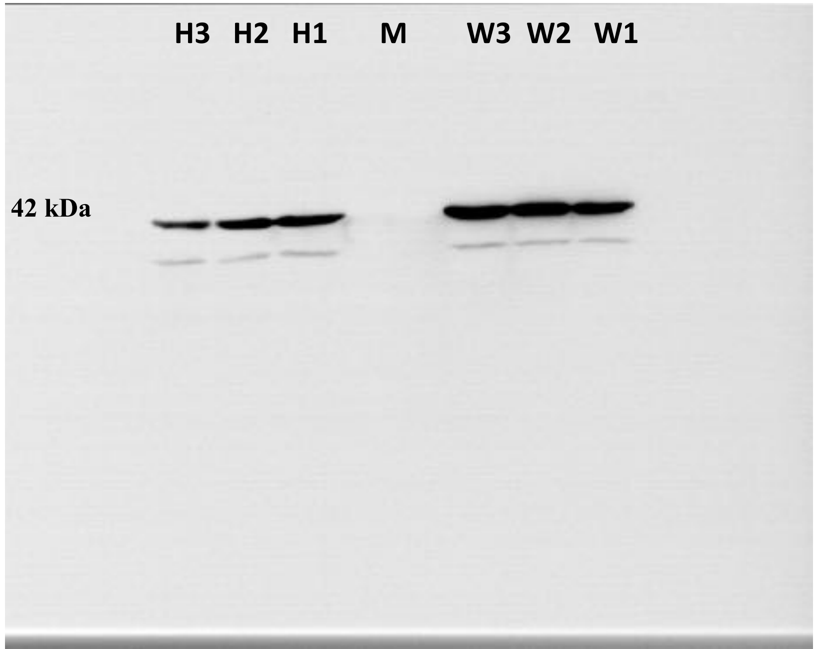

Figure 5B –P28

M H3 H2 H1 W3 W2 W1 M

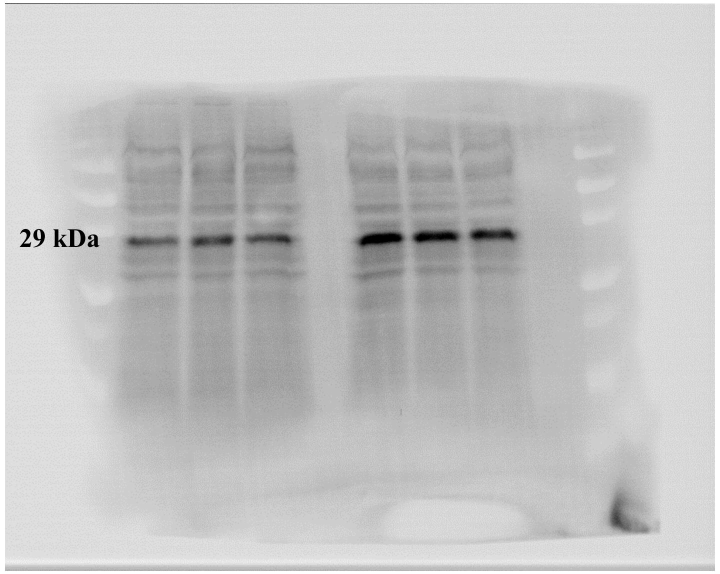


M H3 H2 H1 W3 W2 W1 M

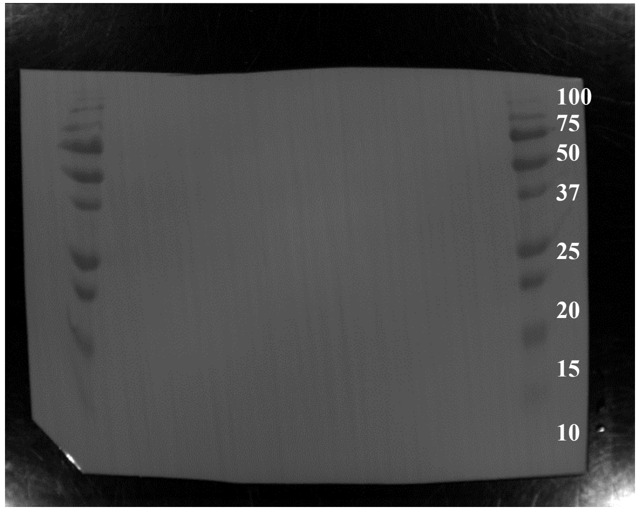


M H3 H2 H1 W3 W2 W1 M

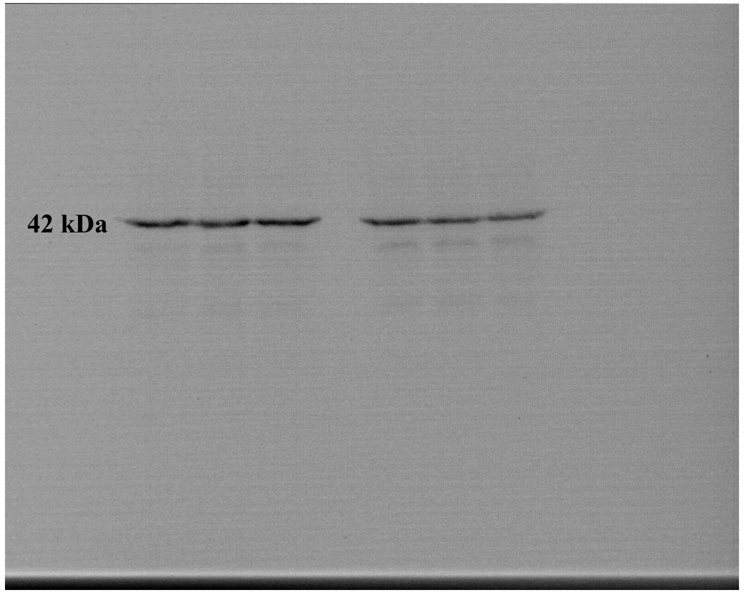


M H3 H2 H1 W3 W2 W1 M

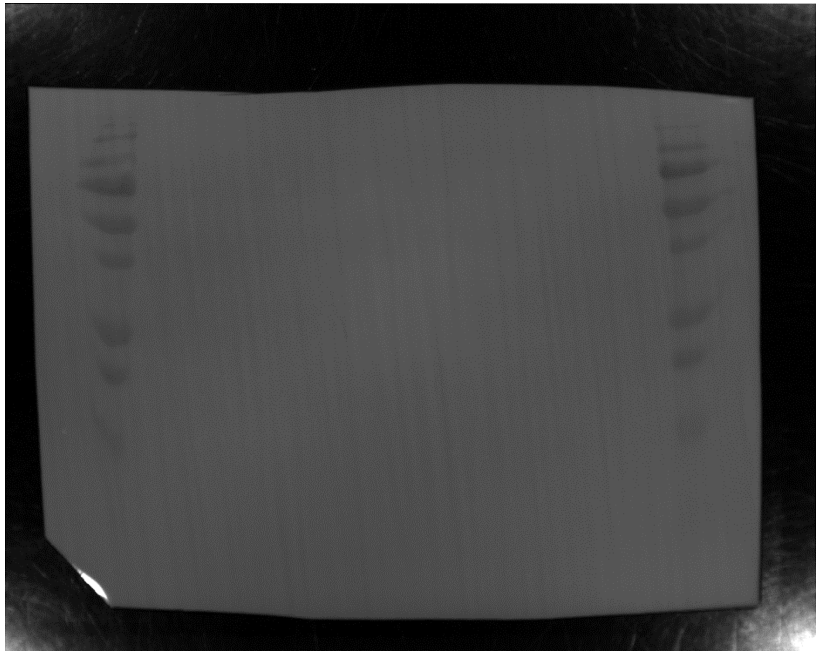

### Figure 5E –P0

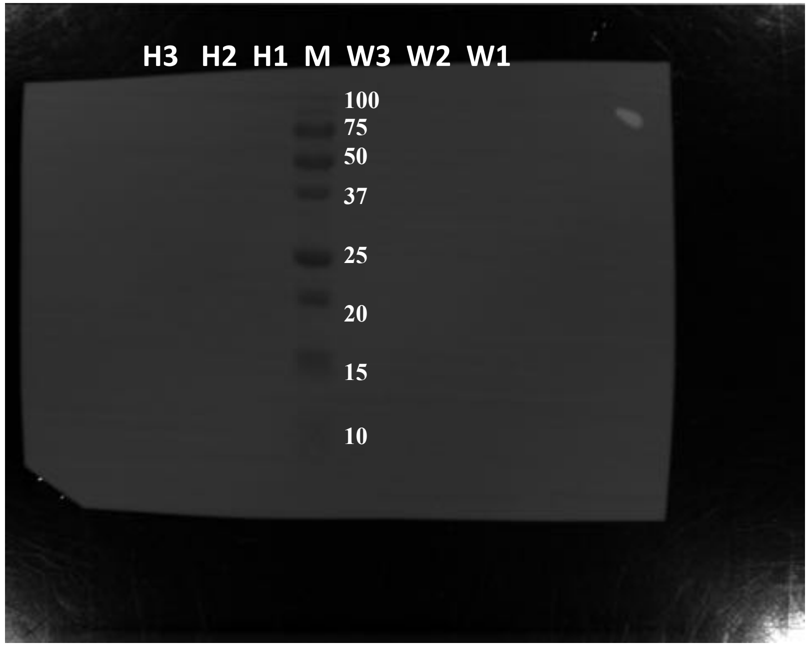
H3 H2 H1 M W3 W2 W1
100
75
50
37
25
20
15
10

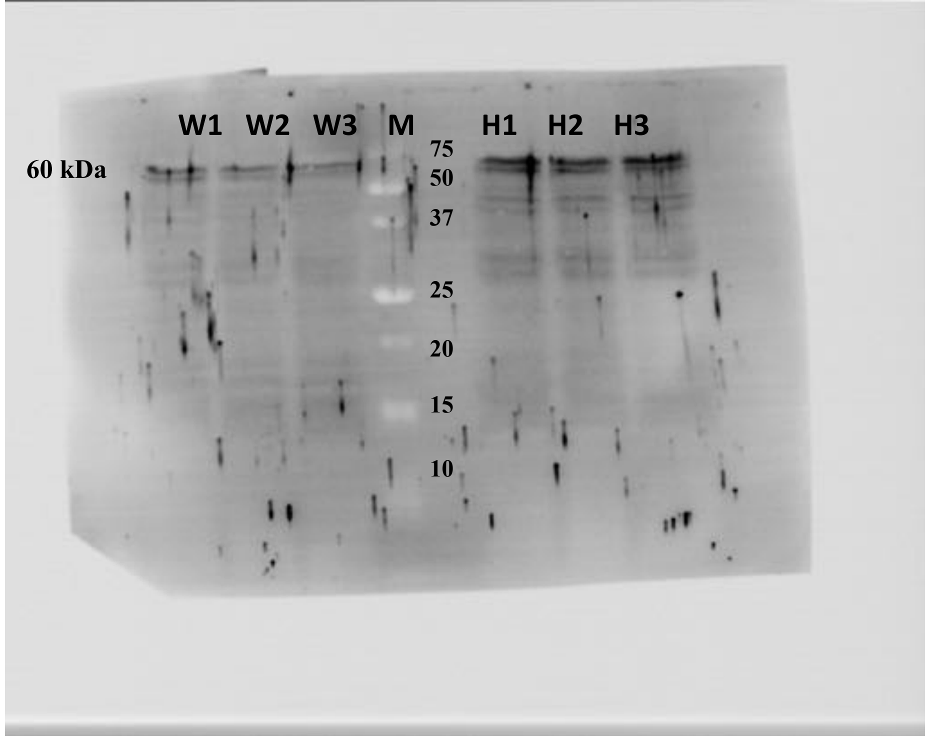
W1 W2 W3 M H1 H2 H3
60 kDa
75
50
37
25
20
15
10

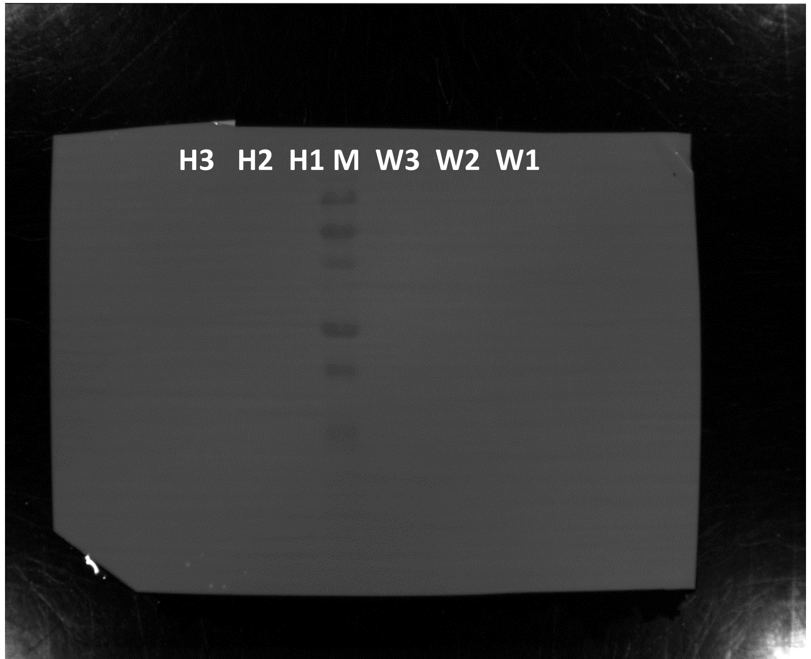
H3 H2 H1 M W3 W2 W1

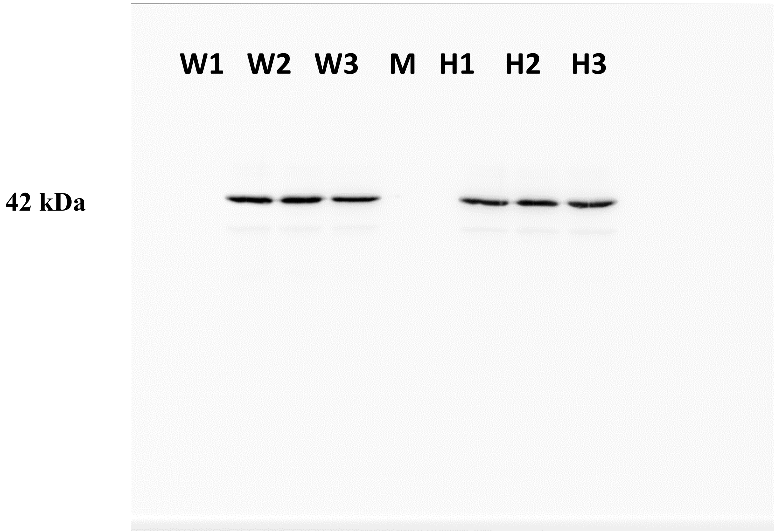
W1 W2 W3 M H1 H2 H3
42 kDa

### Figure 5E –P16

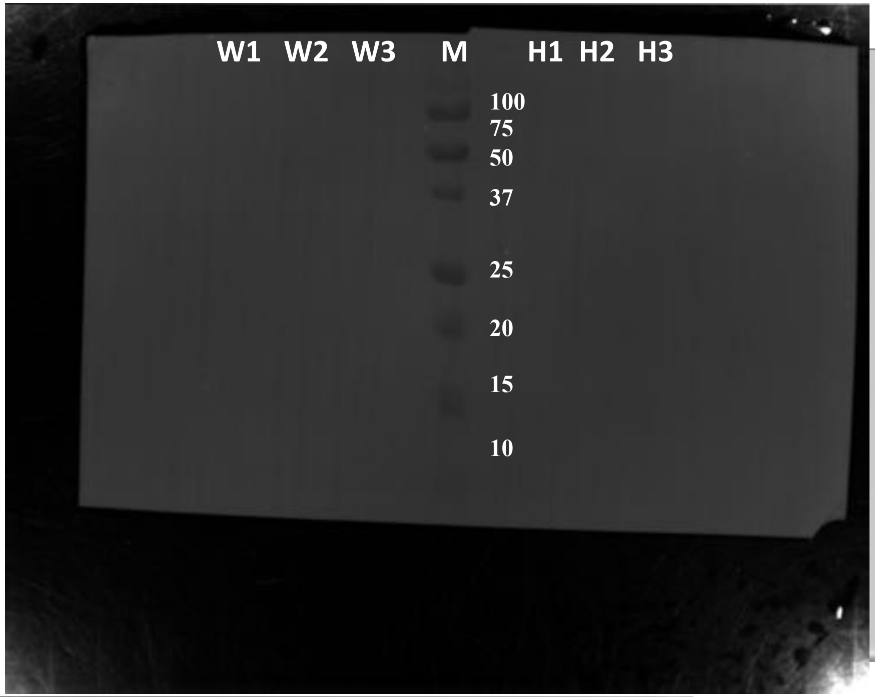


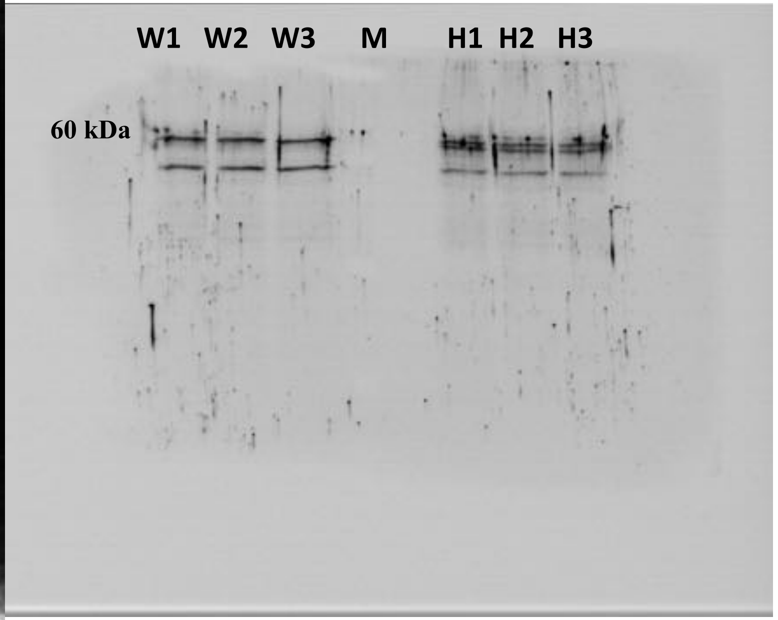


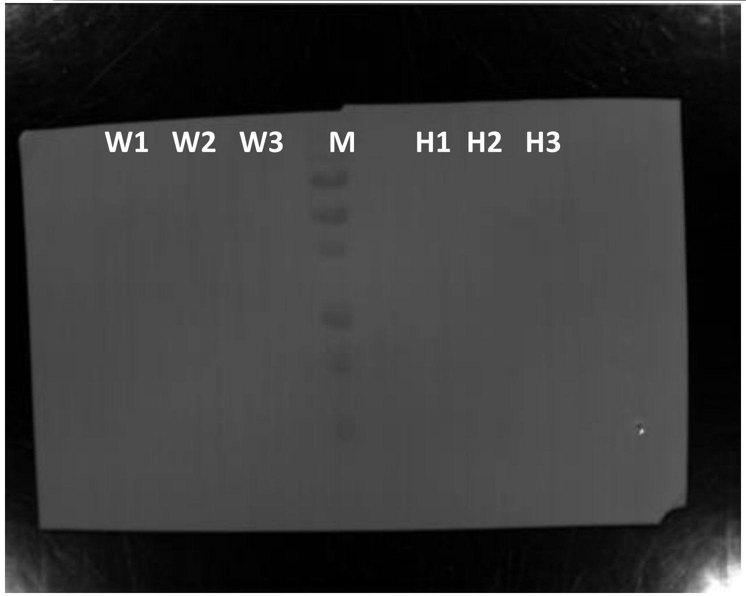


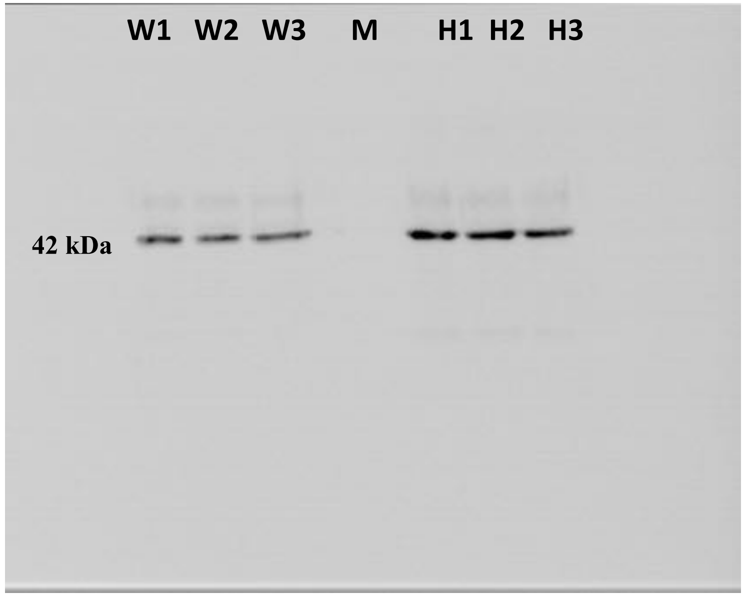

Figure 5E –P19

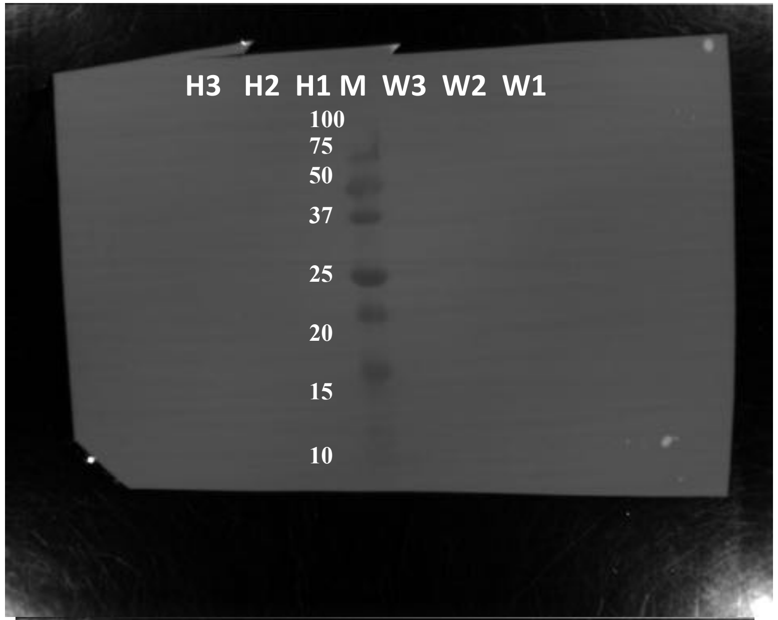


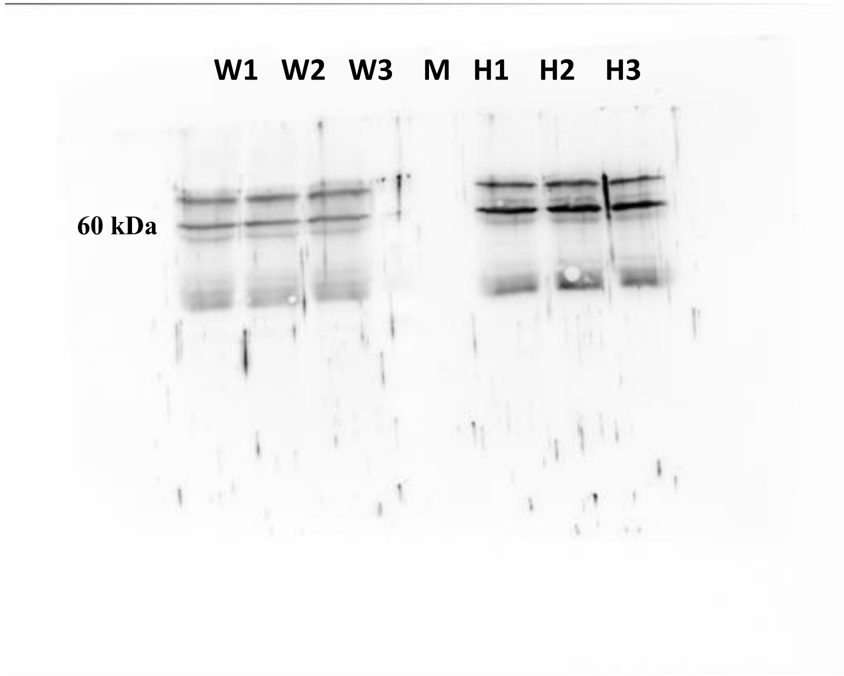


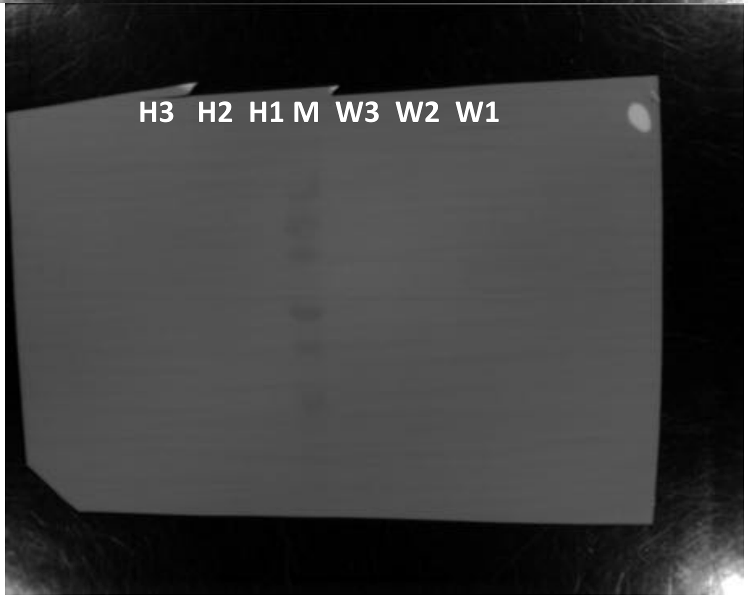


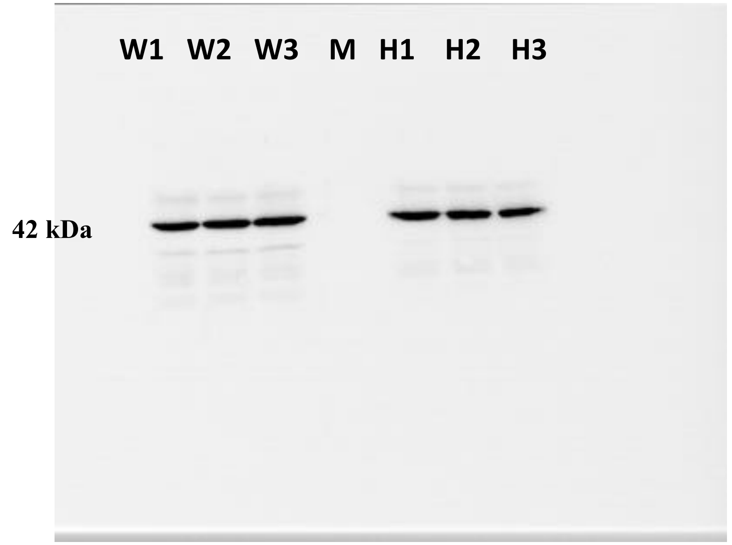

### Figure 5E –P28

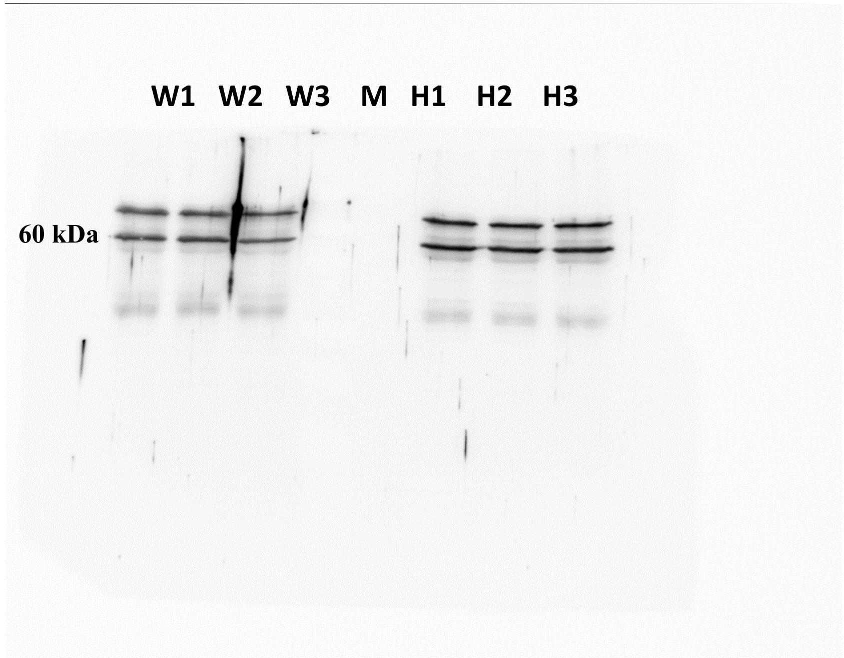


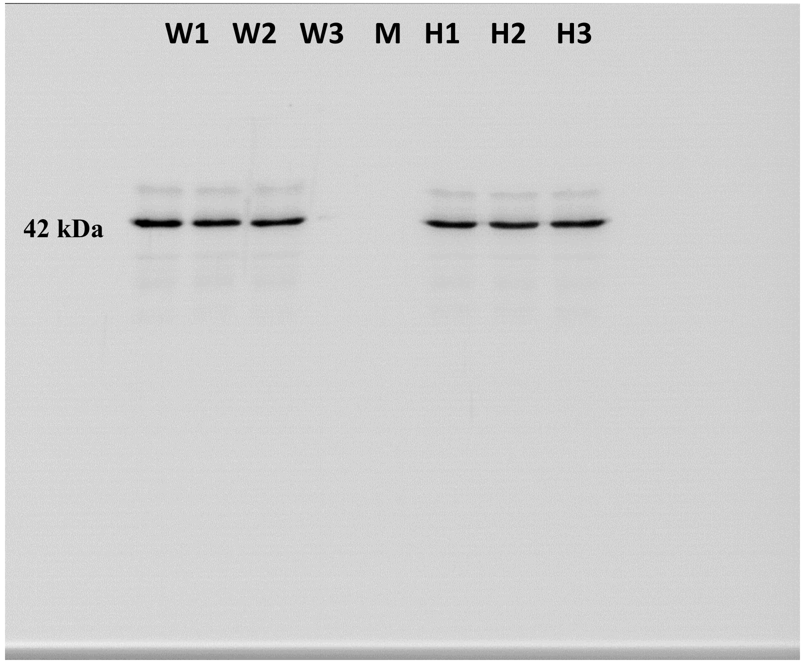


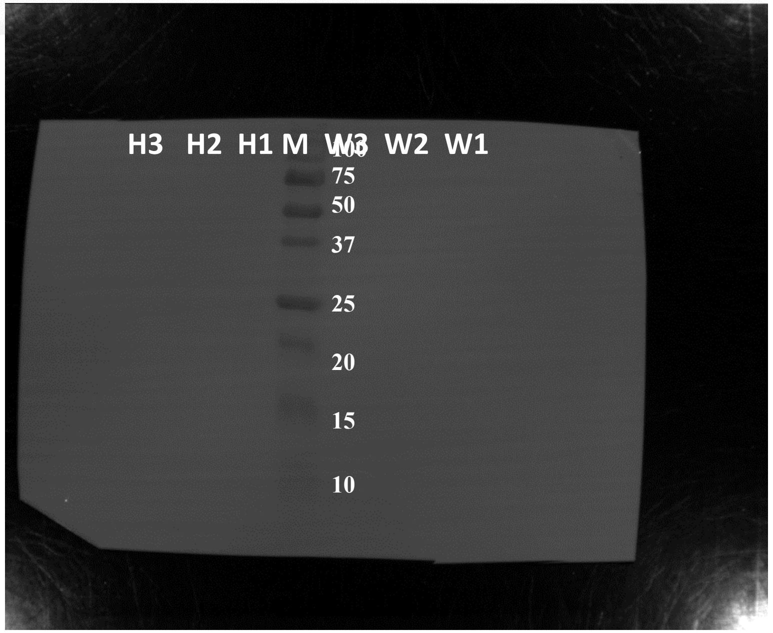


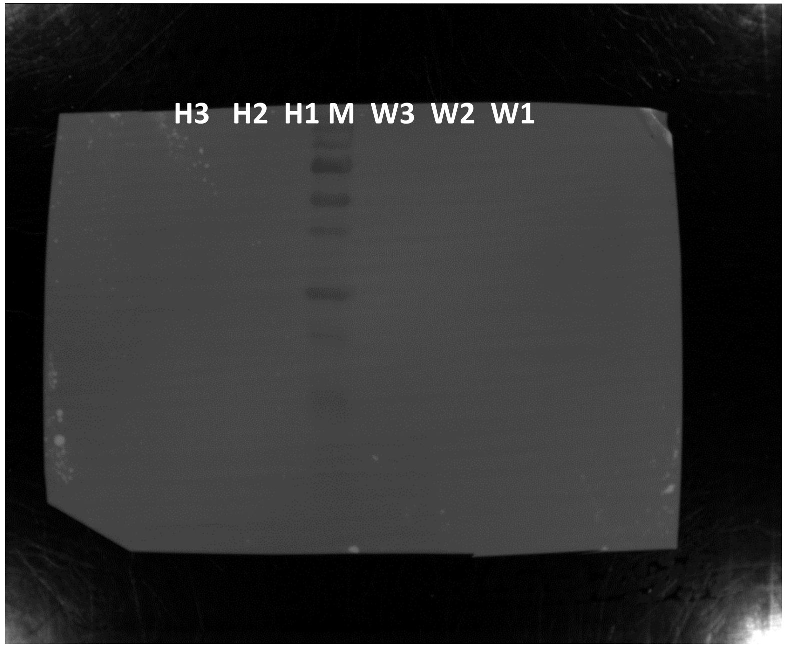
